# Tyrosine kinase inhibitors trigger lysosomal damage-associated cell lysis to activate the NLRP3 inflammasome

**DOI:** 10.1101/2022.02.19.480941

**Authors:** Emilia Neuwirt, Giovanni Magnani, Tamara Ćiković, Anna Kostina, Svenja Wöhrle, Stephan Flemming, Larissa Fischer, Nora J. Fischenich, Benedikt S. Saller, Oliver Gorka, Steffen Renner, Claudia Agarinis, Christian Parker, Andreas Boettcher, Christopher J. Farady, Rolf Backofen, Marta Rodriguez-Franco, Martina Tholen, Thomas Reinheckel, Thomas Ott, Christina J. Groß, Philipp J. Jost, Olaf Groß

## Abstract

Inflammasomes are intracellular protein complexes that control proteolytic maturation and secretion of inflammatory interleukin-1 (IL-1) family cytokines and are thus important in host defense. While some inflammasomes are activated simply by binding to pathogen-derived molecules, others, including those nucleated by NLRP3 and NLRP1, have more complex activation mechanisms that are not fully understood. We screened a library of small molecules to identify new inflammasome activators that might shed light on activation mechanisms. In addition to validating dipeptidyl peptidase (DPP) inhibitors as NLRP1 activators, we find that clinical tyrosine kinase inhibitors (TKIs) including imatinib and masitinib activate the NLRP3 inflammasome. Mechanistically, these TKIs cause lysosomal swelling and damage, leading to cathepsin-mediated destabilization of myeloid cell membranes and cell lysis. This is accompanied by potassium (K^+^) efflux, which activates NLRP3. Both lytic cell death and NLRP3 activation but not lysosomal damage induced by TKIs are prevented by the cytoprotectant high molecular weight polyethylene glycol (PEG). Our study establishes a screening method that can be expanded for inflammasome research and immunostimulatory drug development, and provides new insight into immunological off-targets that may contribute to efficacy or adverse effects of TKIs.

**One Sentence Summary:** A functional small molecule screen identifies imatinib, masitinib and other tyrosine kinase inhibitors that destabilize myeloid cell lysosomes, leading to cell lysis and K^+^ efflux-dependent NLRP3 inflammasome activation.

## Introduction

Inflammasomes are intracellular danger-sensing complexes that couple the detection of pathogen-derived molecules and other danger signals to the secretion of mature IL-1 family cytokines and to a form of lytic cell death termed pyroptosis (*1, 2*). They are typically found in myeloid cells of the innate immune system and are important for host defense against infection, but are also implicated in deleterious inflammatory responses in numerous diseases (*3*). Minimally, they consist of an oligomerized sensor moiety (often a protein of the NOD-like receptor [NLR] family) connected to the protease caspase-1, usually via a large oligomeric complex consisting of the adaptor protein ASC (*4, 5*). Following its activation at the inflammasome, caspase-1 cleaves not only pro-IL-1β and pro-IL-18 to produce the active cytokines, but also the pore-forming protein gasdermin D (GSDMD) that mediates IL-1 release and executes pyroptosis (*6*). Like other lytic cell death modalities such as necroptosis, pyroptosis involves osmotic cell swelling (*7*) and loss of plasma membrane integrity, and also leads to the release of the cellular contents including damage-associated molecular patterns and alarmins that add their immunostimulatory potential to that of IL-1 cytokines (*8*).

Several inflammasomes are activated by direct binding of a pathogen-derived ligand in the cytoplasm: for instance, the NAIP/NLRC4 inflammasomes detects bacterial secretion system components or flagellin (*9*) and the AIM2 inflammasome binds cytoplasmic dsDNA (*10*). Experimentally, precision activation of these inflammasomes can be achieved simply by transfecting the aforementioned ligands. Other inflammasomes sense pathogen activity indirectly and are therefore more challenging to activate specifically. NLRP1 paralogs are hypothesized to detect pathogens by acting as decoy substrates for pathogen-derived enzymes such as proteases or E3 ubiquitin ligases that target NLRP1 for partial proteasomal degradation, thereby relieving the remaining portion from auto-inhibition (*11*). Pyrin (encoded by *Mefv*) is kept inactive by phosphorylation mediated by homeostatic Rho GTPase signalling; it is triggered by pathogen effectors that suppress RhoA activity, leading to its dephosphorylation and activation (*12*).

The NLRP3 inflammasome is unique in that it has a myriad of structurally diverse activators of exogenous and endogenous origin, and in that its precise mechanism of activation is not well understood (*13*). Since its discovery as a gene mutated in hereditary fever syndromes, NLRP3 has been implicated in numerous acquired diseases and inflammatory conditions (*3*). Consistent with its role in inflammatory diseases, NLRP3 is activated by endogenous danger signals produced during tissue damage (*e.g.* extracellular ATP) or metabolic deregulation and excess (*e.g.* saturated fatty acids, or crystals of monosodium urate [MSU], oxalate or cholesterol) (*14*). Furthermore, the ability of NLRP3 to respond to environmental irritants explains its pathogenic role in asbestosis, silicosis and contact hypersensitivity. NLRP3 can also be activated by pore-forming toxins such as nigericin, as well as certain pathogens (*15*) and small molecules (*16*). None of the numerous structurally unrelated NLRP3 activators are known to act as direct ligands, although they have common effects on the cell, which in turn appear to be involved in NLRP3 activation. These include potassium (K^+^) efflux (*17, 18*), metabolic dysregulation and mitochondrial ROS production (*14, 16*), and disturbance of intracellular membranous compartments such as the lysosome (*19, 20*).

Efforts to discover small molecule modulators of the inflammasome have primarily focused on identification of inhibitors (*21–23*), though some have also identified new activators (*24*). Small molecule activators of NLRP1, pyrin and NLRP3 would simplify the study of these inflammasomes and, particularly in the case of NLRP3, might yield fundamental insight into activation mechanisms. For instance, our previous investigation dissecting the mechanism of NLRP3 activation by imiquimod (R837) and related imidazoquinolines revealed that K^+^ efflux is not a universal requirement for NLRP3 activation (*16*). Furthermore, while studies performed in gene-deficient mice have implicated inflammasomes in several inflammatory diseases and in host defense, less is known about the drugs that activate the inflammasome and, by extension, the therapeutic settings in which inflammasomes might play a role in either efficacy or side effects of a given drug. In addition, specific novel small molecule activators of inflammasomes have therapeutic potential as immunostimulatory drugs, for instance in the context of cancer therapy or vaccination since the established inflammasome activators have pleiotropic effects or are not suitable in a therapeutic setting.

We therefore screened a library of small molecules including known drugs for their ability to activate the inflammasome. We identified Val-boro-Pro as an activator of the NLRP1 inflammasome, which is in line with other recent studies (*25–29*). Furthermore, we identified the TKIs imatinib (Glivec™, Gleevec™) and the structurally-related masitinib (Masivet™) as novel activators of the NLRP3 inflammasome. Specifically in myeloid cells, imatinib and masitinib triggered lysosomal swelling and damage, and subsequent plasma membrane destabilization and ballooning, ultimately resulting in cell lysis and K^+^ efflux to activate NLRP3. Stabilizing the cell membrane with the cytoprotectant high molecular weight PEG blocked NLRP3 inflammasome activation by imatinib and masitinib and also by MSU as a crystaline activator of the lysosomal NLRP3 activation pathway (*19, 30*). Other clinically approved antineoplastic TKIs such as bosutinib (Bosulif™) and crizotinib (Xalkori™) also triggered NLRP3 activation in a similar fashion in murine and human myeloid cells, suggesting this may be a common feature of this family of drugs with potential relevance in the therapeutic setting.

## Results

### Small molecule screen for inflammasome activators

To identify new small molecule inflammasome activators, we designed a two-step screening strategy employing loss of cell viability (utilizing a luciferase-based assay which monitors reduction in cellular ATP levels) and IL-1β secretion as primary and secondary end-points, respectively (**Fig. 1A**). The screen was performed in a 384-well format using lipopolysaccharide (LPS)-primed primary murine bone marrow-derived dendritic cells (BMDCs), as they display robust and rapid inflammasome activation (*31, 32*). Nigericin is an established NLRP3 activator and was used as the positive control. We screened a library of 2256 small molecules that included cytostatic agents and signal transduction inhibitors, as well as other established pharmaceuticals, candidate drugs and research compounds for which at least some indication for a mode of action exists (*33*). When we tested at a concentration of 50 µM, 21% of the compounds reduced cell viability by at least 50% (**Fig. 1B**). To identify the inflammasome activators among these compounds, we performed a secondary screen using the same experimental conditions and a FRET-based assay that detects the cleaved and released form of IL-1β (*34*). A total of 98 compounds induced the release of at least 50% of the amount of IL-1β induced by nigericin (**Fig. 1C**). As compared to the composition of the library, we found that compounds targeting proteases and kinases formed the largest groups among the hits. Furthermore, compounds targeting kinases were enriched 2.7-fold amongst the hits as compared to the complete library (**Fig. 1D**). We selected compounds from these categories for in-depth analysis (see **Supplementary Table 1** for structures and trade names of the drugs analyzed throughout this study).

**Fig. 1:**
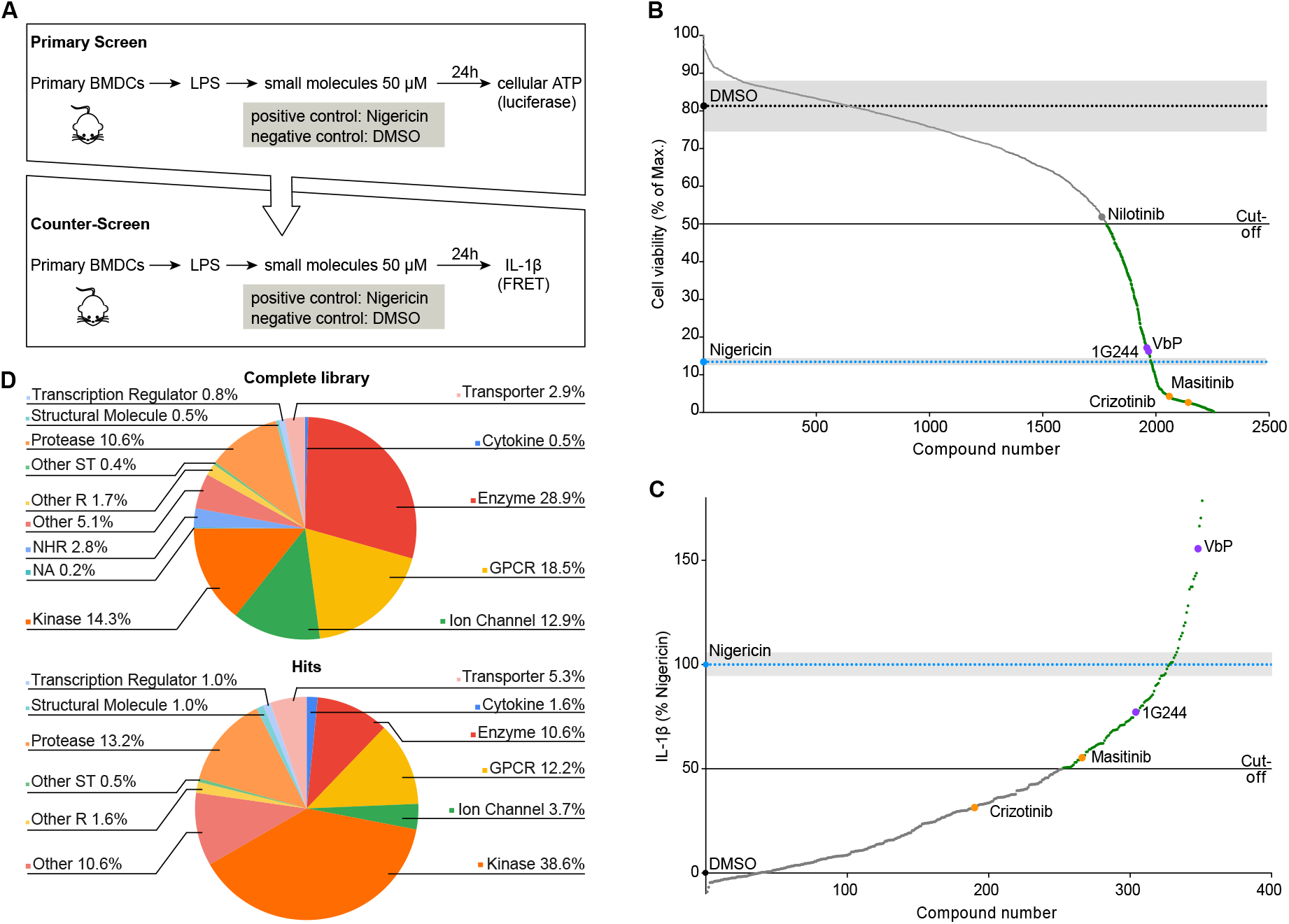
Small molecule screen for inflammasome activators. (**A**) Schematic overview of the two-step screening strategy. (**B**) Cell viability of LPS-primed BMDCs was assessed by determining total cellular ATP after incubation with the compound library at 50 µM for 20 h. 5 µM nigericin was used as a positive or 0.5% DMSO as a negative control for loss of viability. (**C**) BMDCs were primed with LPS and stimulated with compounds identified from the primary screen (B) at 50 µM for 20 h. Release of IL-1β into the supernatants was determined by FRET-based HTRF technology and is depicted as % of IL-1β release induced by 5 µM nigericin. (**D**) Global analysis of the targets of the 98 hits as compared to the whole library. Description of which mode-of-actions / targets / structural characteristics are enriched. GPCR: G protein-coupled receptor, NHR: nuclear hormone receptor, other R: other receptor, other ST: other signal transducer, NA: not available.

### Inflammasome activation by dipeptidyl peptidase inhibitors

Among the strongest inducers of IL-1β secretion identified in the screen was Val-boro-Pro (VbP, also known as talabostat, PT-1000) (**Fig. 1C**), an antineoplastic candidate drug that inhibits dipeptidyl peptidases (DPPs) (*25*). VbP triggered substantial maturation and release of IL-1β, as well as lytic cell death as measured by release of lactate dehydrogenase (LDH) (**Supplementary Fig. 1A-C**). This was accompanied by cleavage of the inflammasome effector protease caspase-1 and its pore-forming substrate GSDMD (**Supplementary Fig. 1C**), as well as formation of ASC ‘specks’ as visualized by immunofluorescence staining and confocal microscopy (**Supplementary Fig. 1D**). Together, these findings suggest the activation of an inflammasome. Similar results were observed with 1G244, another screening hit that inhibits DPP8 and DPP9 with greater specificity than VbP (*25*) (**Fig. 1A-D**).

To gain insight into which inflammasome was activated by the DPP inhibitors, we analyzed cells deficient in ASC (encoded by *Pycard*). Release of mature IL-1β as well as cleavage of caspase-1 were dependent on ASC (**Supplementary Fig. 1A, C**). However, in contrast to what was observed with the control NLRP3 activator nigericin, significant GSDMD processing (**Supplementary Fig. 1C**) and pyroptotic cell death (**Supplementary Fig. 1B**) were still observed in the absence of ASC but not in the absence of caspase-1. This suggested that the DPP inhibitors activate an inflammasome-nucleating receptor such as NLRP1 or NLRC4 that contain a caspase recruitment domain (CARD) and can thereby directly engage caspase-1 for partial activation in the absence of ASC (*1, 35*). In contrast to other inflammasome-nucleating proteins, NLRP1 activation requires its partial degradation by the proteasome, releasing its C-terminus from auto-inhibition (*36*). The proteasome inhibitor MG132 blocked IL-1β release induced by VbP and 1G244, but not by the NLRP3 inflammasome activator nigericin, or the AIM2 inflammasome activator poly(dA:dT), in line with recent reports that DPP inhibitors activate the NLRP1 inflammasome (*25–29*) (**Supplementary Fig. 1E**). Cells from C57BL/6 mice are insensitive to NLRP1 activation by the established stimulus Anthrax lethal toxin that in mice triggers only the NLRP1b paralog absent in this strain (*27*). Therefore, the markedly enhanced sensitivity to VbP observed in cells from BALB/c mice is consistent with NLRP1 activation (**Supplementary Fig. 1F**) but suggests that DPP inhibitors activate NLRP1b and at least one other NLRP1 paralog, which is consistent with other studies (*25–29*). These studies also report that the ability of this class of compounds to activate the inflammasome is related to their inhibition of both DPP8 and DPP9 and a direct DPP interaction with NLRP1, which is consistent with our observation that the DPP4 inhibitor alogliptin did not trigger inflammasome activation (**Supplementary Fig. 1G**). The identification of DPP inhibitors by our screen demonstrates that our screening strategy is suitable for discovery of new small molecule inflammasome activators.

### Imatinib and masitinib activate the NLRP3 inflammasome

Interestingly, the antineoplastic tyrosine kinase inhibitor (TKI) masitinib was among the compounds that most strongly reduced ATP levels (*i.e.* BMDC viability) in the primary screen, and also induced IL-1β release in the secondary screen (**Fig. 1B, C**). We also tested the prototypic TKI imatinib, which is structurally similar to masitinib (**Supplementary Table 1**), and found that both compounds induced IL-1β and also triggered caspase-1 cleavage and formation of ASC specks (**Fig. 2A, B** and **Supplementary Fig. 2A-E**). The approximate minimum dose and duration required for strong IL-1β secretion (*i.e.* comparable to established inflammasome activators) were 20 µM and 2-3 h, respectively (**Supplementary Fig. 2A, B**). Release of mature IL-1β and cleavage of caspase-1 and GSDMD in response to imatinib and masitinib were strongly reduced in the absence of NLRP3 in BMDCs (**Fig. 2A, B** and **Supplementary Fig. 2E**). Similarly, imatinib and masitinib induced robust, NLRP3-dependent IL-1β production in human PMA (phorbol 12-myristate 13-acetate)-differentiated THP-1 cells (myeloid cell line) and in LPS-primed murine bone marrow-derived macrophages (BMDMs) (**Fig. 2C** and **Supplementary Fig. 2F**), suggesting that imatinib and masitinib are novel activators of the NLRP3 inflammasome in murine and human myeloid cells. Indeed, MCC950, a small molecule that directly inhibits NLRP3 (*37, 38*), also inhibited IL-1β secretion in response to imatinib, masitinib and other NLRP3 activators both in murine BMDCs and LPS-primed human peripheral blood mononuclear cells (PBMCs), without interfering with AIM2 inflammasome activation by poly(dA:dT) (**Fig. 2D, E**).

**Fig. 2:**
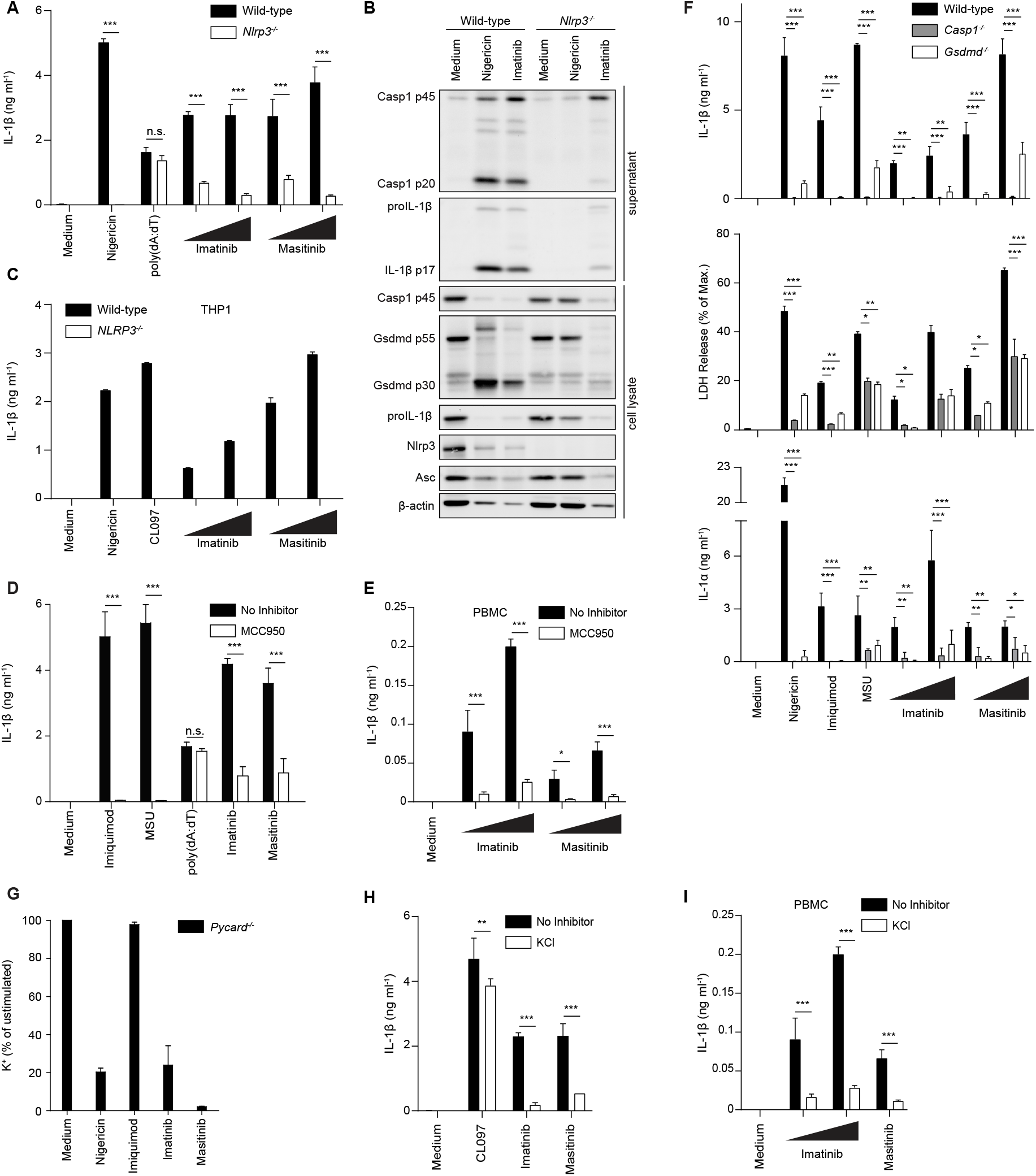
Imatinib and masitinib activate the NLRP3 inflammasome and trigger inflammasome-independent lytic cell death. (**A**) BMDCs from wild-type and *Nlrp3^-/-^*mice were primed with LPS and then stimulated for 3 h with 5 µM nigericin, 2 µg ml^-1^ poly(dA:dT), 40 µM and 60 µM imatinib, or 20 µM and 40 µM masitinib, respectively and IL-1β secretion was measured. (**B**) Immunoblot analysis of cell lysates and supernatants from wild-type and NLRP3-deficient, LPS-primed BMDCs stimulated with 5 µM nigericin, 40 µM imatinib, or left untreated for 3 h. (**C**) Wild-type and *NLRP3^-/-^*THP-1 cells were treated with 200 ng ml^-1^ PMA for 3 h, washed and left at 37°C, 5% CO_2_ overnight. The cells were subsequently primed with 150 ng ml^-1^ LPS for 3 h and treated with 5 µM nigericin, 100 µM CL097 or 60 µM and 80 µM of the respective TKI for 3 h and IL-1β secretion was measured. Values in NLRP3-deficient cells were below detection limit. (**D**) LPS-primed BMDCs were treated with 5 µM MCC950 30 min prior to stimulation with 100 µM imiquimod, 300 µg ml^-1^ MSU, 2 µg ml^-1^ poly(dA:dT) and 40 µM imatinib or masitinib and IL-1β secretion was measured. (**E**) LPS-primed human PBMCs were treated with 3 µM MCC950 30 min prior to stimulation with 60 µM and 80 µM imatinib or masitinib for 3 h and IL-1β secretion was measured. (**F**) LPS-primed BMDCs from wild-type, *Casp1^-/-^ and Gsdmd^-/-^* mice were stimulated with 5 µM nigericin, 100 µM imiquimod, 300 µg ml^-1^ MSU or increasing concentrations of imatinib (40 µM, 60 µM) and masitinib (20 µM, 40 µM) and IL-1 secretion and LDH release were measured. (**G**) LPS-primed BMDCs from ASC-deficient *Pycard^-/-^* mice were stimulated with 5 µM nigericin, 100 µM imiquimod, 60 µM imatinib, or 40 µM masitinib. Intracellular K^+^ concentrations were determined by total reflection x-ray fluorescence analysis (TXRF). Data is depicted as percentage K^+^ content of unstimulated cells. (**H**) LPS-primed BMDCs were incubated with 60 mM KCl in BMDC medium or BMDC medium for 30 min and then treated with 100 µM CL097, 40 µM imatinib or 20 µM masitinib for 3 h and IL-1β secretion was measured. (**I**) LPS-primed human PBMCs were incubated with 60 mM KCl 30 min prior to stimulation with 60 µM and 80 µM imatinib or 80 µM masitinib and IL-1β secretion was measured. Cytokine secretion and LDH release were determined by ELISA or using a colorimetric assay, respectively, from cell-free supernatants and data are depicted as mean ±SD of technical triplicates. Results are representative of at least three independent experiments. Multiple unpaired T-tests were performed for statistical analysis (*, p < 0.05; **, p < 0.01; ***, p < 0.001; n.s., not significant).

### Imatinib and masitinib trigger inflammasome-independent lytic cell death and K^+^ efflux- dependent NLRP3 activation

Interestingly, while imatinib- or masitinib-induced secretion of mature IL-1β was completely dependent on caspase-1 (**Fig. 2F**), these TKIs induced significant cell death even in cells lacking caspase-1 or GSDMD, indicating that they induce another lytic cell death mechanism in addition to caspase-1/GSDMD-dependent pyroptosis (**Fig. 2F**). Consistent with this, LDH release was also largely independent of NLRP3 and ASC and also occurred in the absence of LPS as a priming stimulus (**Supplementary Fig. 3A-C**), confirming that TKI can trigger cell death independent of the canonical NLRP3 inflammasome. Furthermore, inflammasome- and priming-independent release of alarmins such as IL-1α or HMGB1 (**Fig. 2F** **and Supplementary Fig. 3D**) and priming-independent membrane damage (measured by Draq7 uptake) and inflammasome activation (determined by ASC speck formation) (**Supplementary Fig. 3E**) demonstrate the strong and multifaceted proinflammatory potential of these TKIs due to their ability to trigger cell death, as well as their ability to trigger inflammasome activation even in unprimed cells.

The data to this point indicated that imatinib and masitinib cause an inflammasome-independent form of lytic cell death, while IL-1β maturation in response to TKIs is inflammasome dependent. These observations are reminiscent of previous studies showing that different lytic cell death pathways can engage the NLRP3 inflammasome. Specifically, necroptotic cell death (*39*) or activation of the cytoplasmic LPS sensor caspase-11 (*40, 41*) lead to NLRP3 activation in myeloid cells by causing formation of MLKL or GSDMD membrane pores, respectively, and thereby efflux of K^+^. Extracellular KCl is an established means of blocking K^+^ efflux and NLRP3 activation in response to cell death inducers and also classical NLRP3 activators like nigericin, ATP and MSU (*16*). We therefore hypothesized that TKIs might also engage NLRP3 by causing K^+^ efflux downstream of membrane destabilization/lysis. Indeed, in contrast to imidazoquinolines like imiquimod and CL097, which cause K^+^ efflux- independent NLRP3 activation (*16*), imatinib and masitinib triggered K^+^ efflux and also required K^+^ efflux for NLRP3 activation (**Fig. 2G-I**). By contrast, IL-1β secretion induced by imatinib and masitinib was intact in cells deficient in the necroptosome protein RIPK3, the non-canonical inflammasome protein caspase-11, or the pyroptotic pore protein GSDMD (**Supplementary Fig. 4A, B and** **Fig. 2F**). Other lytic cell modalities that might in principle also engage NLRP3 via K^+^ efflux are ferroptosis, parthanatos and pyroptosis mediated by another gasdermin family member expressed in these cells, GSDME (*42*). Inhibitors of ferroptosis and parthanatos did not block imatinib and masitinib-induced LDH release (**Supplementary Fig. 4C and D**). Cells from GSDME-deficient mice likewise displayed an intact TKI-induced cell lysis or IL-1β secretion (**Supplementary Fig. 4E**). Collectively, these observations indicate that numerous established programmed lytic cell death modalities do not account for NLRP3 activation by imatinib and masitinib.

### Imatinib and masitinib induce myeloid cell-specific lytic cell death

We postulated that the cell death mechanisms induced by imatinib and masitinib are involved in NLRP3 activation, and therefore characterized the death induced by these TKIs in detail. To distinguish these NLRP3-independent upstream mechanisms from downstream effects of inflammasome activation (which also include further lysis via pyroptosis), for many experiments we utilized myeloid cells from inflammasome-deficient mice such as ASC knockouts (*Pycard^-/-^*). Notably, imatinib and masitinib only killed myeloid cells such as BMDCs and macrophages (**Fig. 2** **and Supplementary Fig. 3**), but not other primary cells such as thymocytes or murine embryonic fibroblasts (MEFs), nor various cell lines (HeLa, HEK 293T, NIH-3T3, HCT 116) (**Fig. 3A, B** and **Supplementary Fig. 5A**). Furthermore, BCR-ABL1–dependent K562 cells did not display any cell lysis or LDH release after 3-h exposure to TKIs, and showed only minor cell lysis after 24-h exposure to TKIs (**Supplementary Fig. 5B**). Therefore, there is a degree of specificity that potentially implies regulation or programming in the cell death induced by imatinib and masitinib, and/or fundamental differences in the biology of myeloid cells that make them acutely sensitive to TKI-induced death. If TKIs simply caused general cytotoxicity for instance due to high concentrations, we would expect that these TKIs would also kill other cell types indiscriminately.

**Fig. 3:**
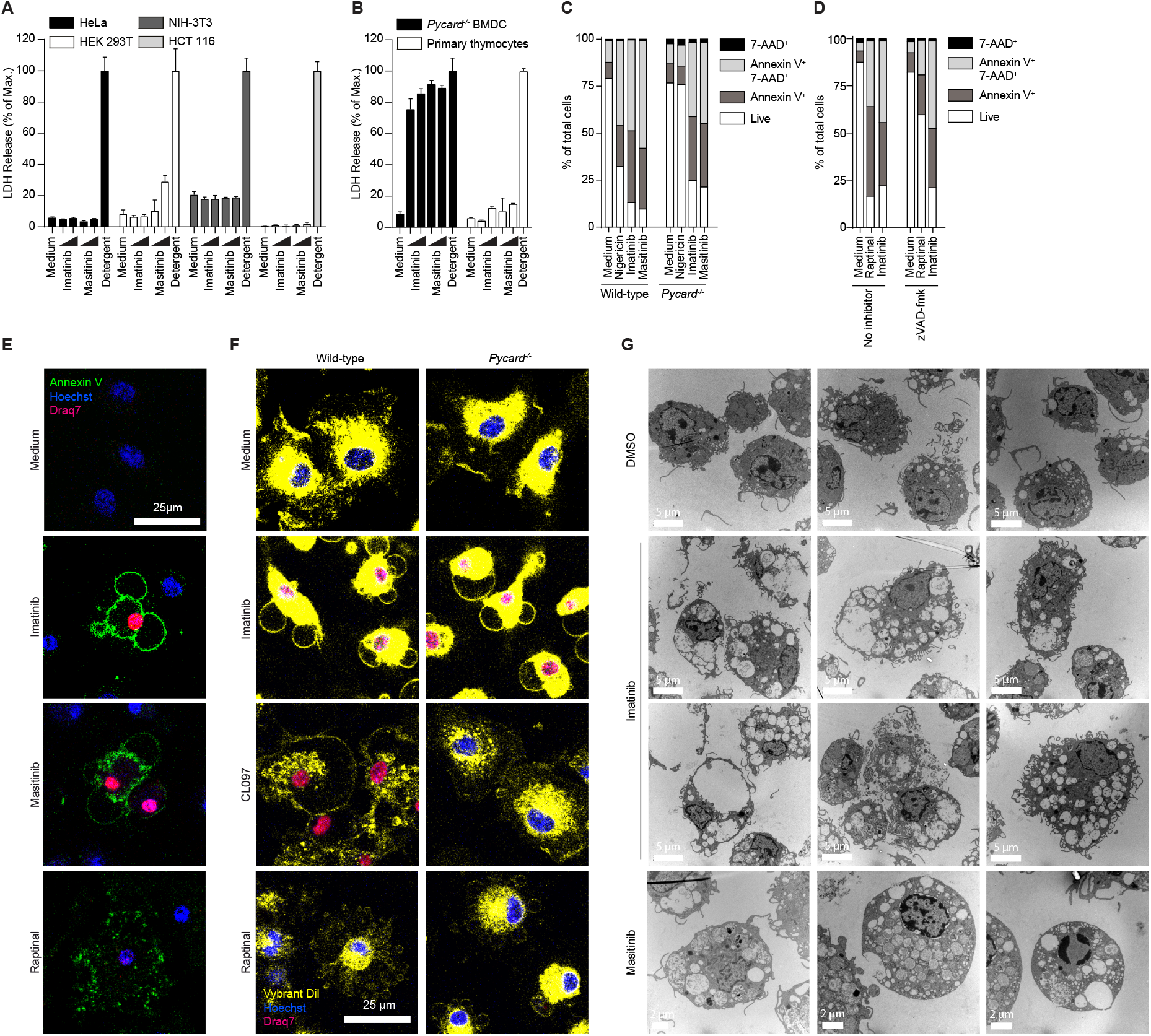
Imatinib and masitinib induce lytic death specifically in myeloid cells. (**A**) and (**B**) HeLa, HEK 293T, NIH-3T3, and HCT 116 cells (A) were treated with 20 and 40 µM imatinib, 10 and 20 µM masitinib or left untreated for 5 h. BMDCs and primary thymocytes from *Pycard^-/-^* mice (B) were incubated with 40 and 60 µM imatinib and masitinib or left untreated for 4 h. Cells were lysed as a positive control for maximum lysis (“Detergent”). LDH was determined from cell-free supernatants by a colorimetric assay and data are depicted as mean (SD) of technical triplicates. (**C**) and (**D**) LPS-primed BMDCs from wild-type and ASC-deficient *Pycard^-/-^*mice were stimulated with 5 µM nigericin, 60 µM imatinib or 40 µM masitinib (C) or NLRP3-deficient BMDCs incubated with 30 µM zVAD-fmk or medium 30 min prior to stimulation with 10 µM raptinal and 40 µM imatinib (D) and the cells were subsequently labelled with Annexin-V/ 7- AAD and analyzed by flow cytometry. (**E**) LPS-primed wild-type BMDMs were stimulated with 40 µM imatinib, 20 µM masitinib, 10 µM raptinal, or left untreated. The cells were stained with FITC Annexin V (green), nuclei were localized with Hoechst 33342 (blue) and DRAQ7 (red) and subsequently imaged by confocal microscopy 30 min after the stimulation. (**F**) LPS-primed wild-type and ASC-deficient *Pycard^-/-^* BMDMs were stimulated with 40 µM imatinib, 100 µM CL097 and 10 µM raptinal. The cell membrane was stained with Vybrant Dil Cell-Labeling Solution (yellow), nuclei were localized with Hoechst 33342 (blue) and DRAQ7 (red). Images were taken before or 60 min after the stimulation by confocal microscopy. (**G**) LPS-primed ASC-deficient *Pycard^-/-^* BMDMs were stimulated with 40 µM imatinib, 20 µM masitinib or DMSO for 2 h. Cells were fixed by adding 1% glutaraldehyde and subsequently analyzed by transmission electron microscopy.

We next characterized the cell death induced by imatinib or masitinib by flow cytometry and microscopy using stains for phosphatidylserine (PS) exposure (Annexin V) and cellular permeability/lysis (7-AAD, Draq7). The TKIs induced exposure of PS and permeability to 7- AAD, and this occurred independently of ASC, suggesting that these events were not a result of pyroptosis as a consequence of canonical NLRP3 inflammasome activation (**Fig. 3C**). Although PS exposure is classically used as a marker of apoptosis, it is not considered specific as it also occurs during other cell death processes (*43*). Furthermore, the pan-caspase inhibitor zVAD-fmk blocked apoptosis induced by raptinal (*44*) but did not prevent PS exposure or 7-AAD uptake in TKI-treated cells, indicating that TKI-induced myeloid cell death is not caused by caspase activity and is therefore unlikely to be apoptosis (**Fig. 3D**).

Imatinib and masitinib induced striking morphological changes in myeloid cells, in particular the formation of large membrane distensions we termed ‘balloons’ to distinguish them from the small, numerous blebs typical of apoptotic cells (**Fig. 3E, F,** and **Supplementary Fig. 5D**). Transmission electron microscopy confirmed large balloons of the cell membrane, as well as swelling of internal membranous organelles and vacuolization in TKI-treated cells (**Fig. 3G, Supplementary Fig. 6**). The morphological changes occurred independently of ASC and were clearly distinct from pyroptosis induced by nigericin or the imiquimod analogue and NLRP3 activator CL097 or apoptosis induced by raptinal (**Fig. 3E-G, Supplementary Fig. 6**). The imatinib-induced changes in membrane morphology and loss of membrane integrity (as indicated by permeability to Draq7) preceded PS exposure (**Supplementary Fig. 5C**). Together with the insensitivity to caspase inhibition (**Fig. 3D**) and the substantial LDH release (*e.g.* **Fig. 2F, 3B**), this shows that the cell death modality induced by TKIs is clearly distinct from apoptosis and that changes in accessibility of PS to Annexin V in TKI-treated cells is likely a consequence of cell lysis rather than the flipping of PS to the outer leaflet, as is characteristic of apoptosis.

### PEG blocks TKI-driven cell lysis and inflammasome activation

To determine whether loss of membrane integrity is an upstream event required for NLRP3 activation in response to TKIs, we examined the effect of the cytoprotectant polyethylene glycol (PEG) in ASC-deficient cells. High molecular weight PEG protects mammalian cells from diverse lytic cell death modalities by preserving membrane integrity (*7, 45*). Preincubation with PEG protected the cells from TKI-induced loss of membrane integrity, exposure of PS, and formation of membrane balloons (**Fig. 4A, B**), but as expected did not prevent raptinal-induced apoptosis (**Fig. 4A**). Furthermore, treatment with either PEG-600 or PEG-3000 prevented TKI-induced cellular ATP loss and LDH release (**Fig. 4C, D**), suggesting that loss of cellular ATP is not the cause but rather a consequence of death induced by TKIs. In contrast, PEG did not interfere with inhibition of tyrosine phosphorylation in BCR-ABL1– dependent K-562 myelogenous leukemia cells (**Supplementary Fig. 7A**) indicating that TKI entry into cells and action on target kinases is intact in the presence of PEG. Furthermore, BMDCs treated with PEG continued to secrete the cytokine tumor necrosis factor (TNF) in response to LPS (**Supplementary Fig. 7B**), indicating that PEG does not prevent basic cellular functions such as protein production and secretion.

**Fig. 4:**
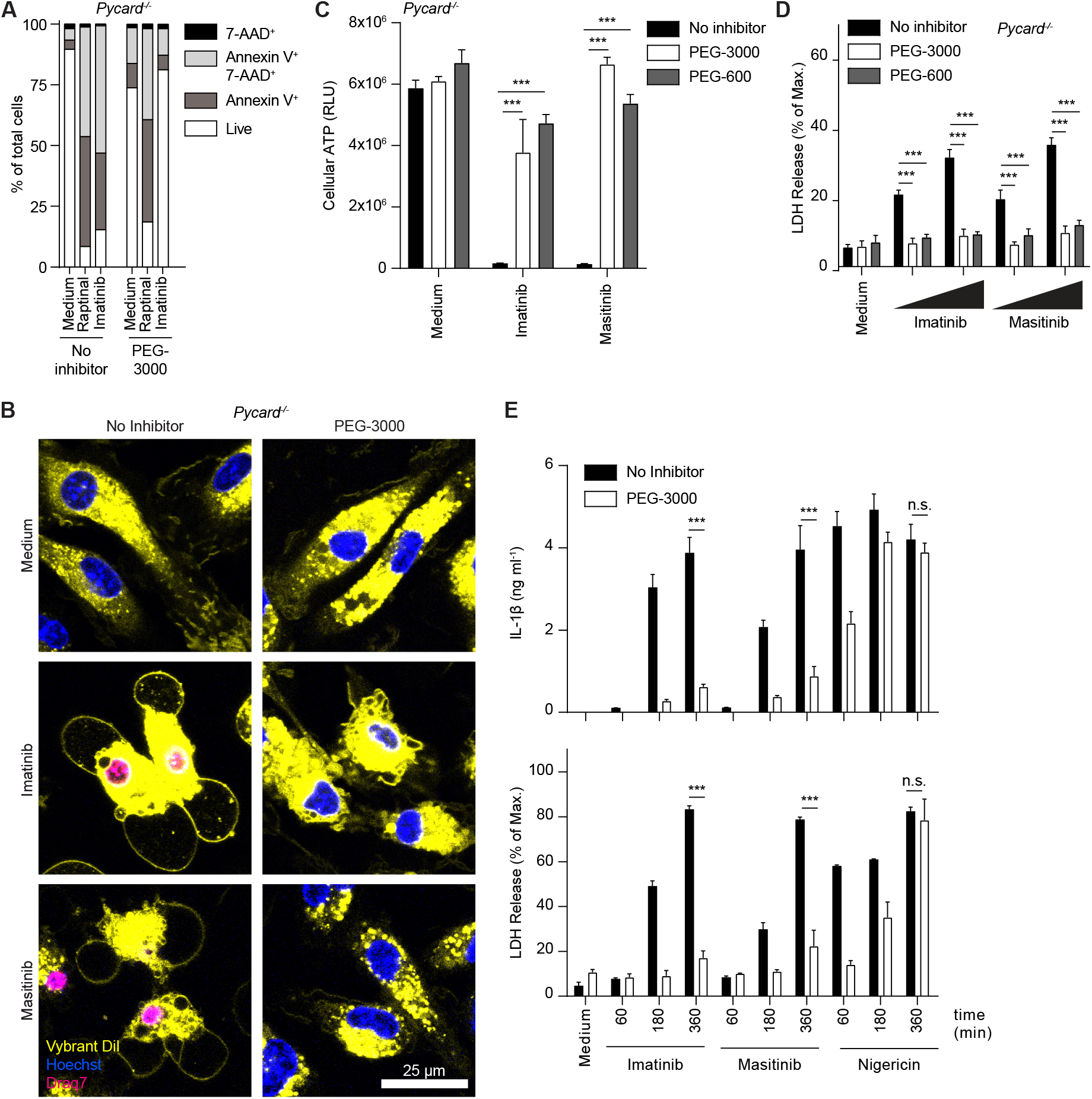
PEG rescues TKI-driven cell lysis and inflammasome activation. (**A**) ASC-deficient *Pycard^-/-^* BMDCs were primed with LPS and then incubated with 10 mM PEG-3000 or left untreated 30 min prior to stimulation with 10 µM raptinal or 40 µM imatinib for 3 h. Cells were stained with Annexin-V and 7-AAD and analyzed by flow cytometry. (**B**) LPS-primed BMDMs from ASC-deficient *Pycard^-/-^* mice were treated with 10 mM PEG- 3000 for 30 min and then stimulated with 40 µM imatinib, 20 µM masitinib or left untreated. The cell membrane was stained with Vybrant DiI Cell-Labeling Solution (yellow), nuclei were localized with Hoechst 33342 (blue) and DRAQ7 (red). Images were taken 60 min after the stimulation. (**C**) and (**D**) LPS-primed ASC-deficient *Pycard^-/-^* BMDCs were incubated with 15 mM PEG- 3000, 150 mM PEG-600 or left untreated for 30 min and then stimulated with 40 µM imatinib and 20 µM masitinib for 3 h. Cellular ATP levels were determined using a luminescent assay (C). (**E**) LPS-primed wild-type BMDCs were incubated with 15 mM PEG-3000 or medium for 30 min. Cells were then stimulated with 40 µM imatinib, 20 µM masitinib or 5 µM nigericin as indicated. IL-1β secretion and LDH release were determined by ELISA or using a colorimetric assay, respectively, from cell-free supernatants and data are depicted as mean ±SD of technical triplicates. Results are representative of at least three independent experiments. Multiple unpaired T-tests were performed for statistical analysis (*, p < 0.05; **, p < 0.01; ***, p < 0.001; n.s., not significant).

Remarkably, PEG blocked not only TKI-induced inflammasome-independent lytic cell death, but also caspase-1 activation and IL-1β secretion, whereas smaller osmoprotectants such as sucrose and glycine did not (**Fig. 4E** and **Supplementary Fig. 7C-E**). In contrast, PEG only slightly delayed NLRP3 inflammasome-dependent pyroptosis and IL-1β secretion induced by nigericin (**Fig. 4E**). This indicates that PEG potently inhibits the TKI-induced inflammasome-independent lytic cell death, but not other types of lytic cell death such as pyroptosis. Importantly, these findings demonstrate that TKI-induced NLRP3 activation requires lytic cell death or membrane destabilization.

### Imatinib and masitinib induce lysosomal membrane destabilization

A plausible hypothesis is that the observed organelle swelling and vacuolization (**Fig. 3** and **Supplementary Fig. 5 and 6**) could be an upstream event of cell lysis. The cellular morphology of TKI-treated myeloid cells observed in transmission electron microscopy (TEM) (**Fig. 3G** and **Supplementary Fig. 6**) is reminiscent of that induced by LLOMe, a soluble peptide that triggers lysosomal damage as well as NLRP3 activation (*46*). MSU crystals and other particulate activators, after partial phagocytic uptake, disrupt lysosomal and cellular membranes and trigger NLRP3 in a K^+^ efflux-dependent manner (**Supplementary Fig. 8A**) (*16–19, 30*). Particulate and LLOMe-induced cell lysis was reported to be inflammasome independent (*8, 47*), which is similar to our observations with TKIs. No precipitate was observed microscopically in TKI-treated conditions (not shown), and the phagocytosis inhibitor cytochalasin D blocked inflammasome activation in response to MSU crystals but not in response to imatinib or masitinib (**Supplementary Fig. 8B**), suggesting that the TKIs do not activating NLRP3 by forming particulates / precipitates. However, imatinib and masitinib nonetheless triggered rapid and substantial lysosomal swelling, as shown by live cell imaging with acridine orange staining (**Fig. 5A, Supplementary Fig. 8C**). PEG pretreatment had no influence on lysosomal swelling itself, suggesting that PEG uncouples lysosomal damage from cell death and NLRP3 activation. Even in the presence of PEG-3000 (to prevent the confounding effects of cell death) TKI-induced lysosomal swelling was followed by lysosomal leakage, as indicated by a loss in the red lysosomal signal of this dye (*19*) (**Fig. 5B**). Lysosomal membrane destabilization is thought to connect to cell lysis and K^+^ efflux for NLRP3 activation through the activity of lysosomal cathepsins in the cytoplasm, which can be blocked by inhibitors such as CA-074Me (*19, 47*). CA-074Me significantly reduced LDH release and IL- 1β secretion in response to MSU and TKIs, while having no effect on NLRP3 inflammasome activation by nigericin (**Fig. 5C**). PEG also prevented MSU particle-induced lysis and IL-1β release (**Fig. 5D**). Together, these results establish high molecular weight PEG as a robust inhibitor of lysosomal membrane permeabilization-induced cell lysis and NLRP3 activation and demonstrate that TKIs activate NLRP3 via the lysosomal pathway.

**Fig. 5:**
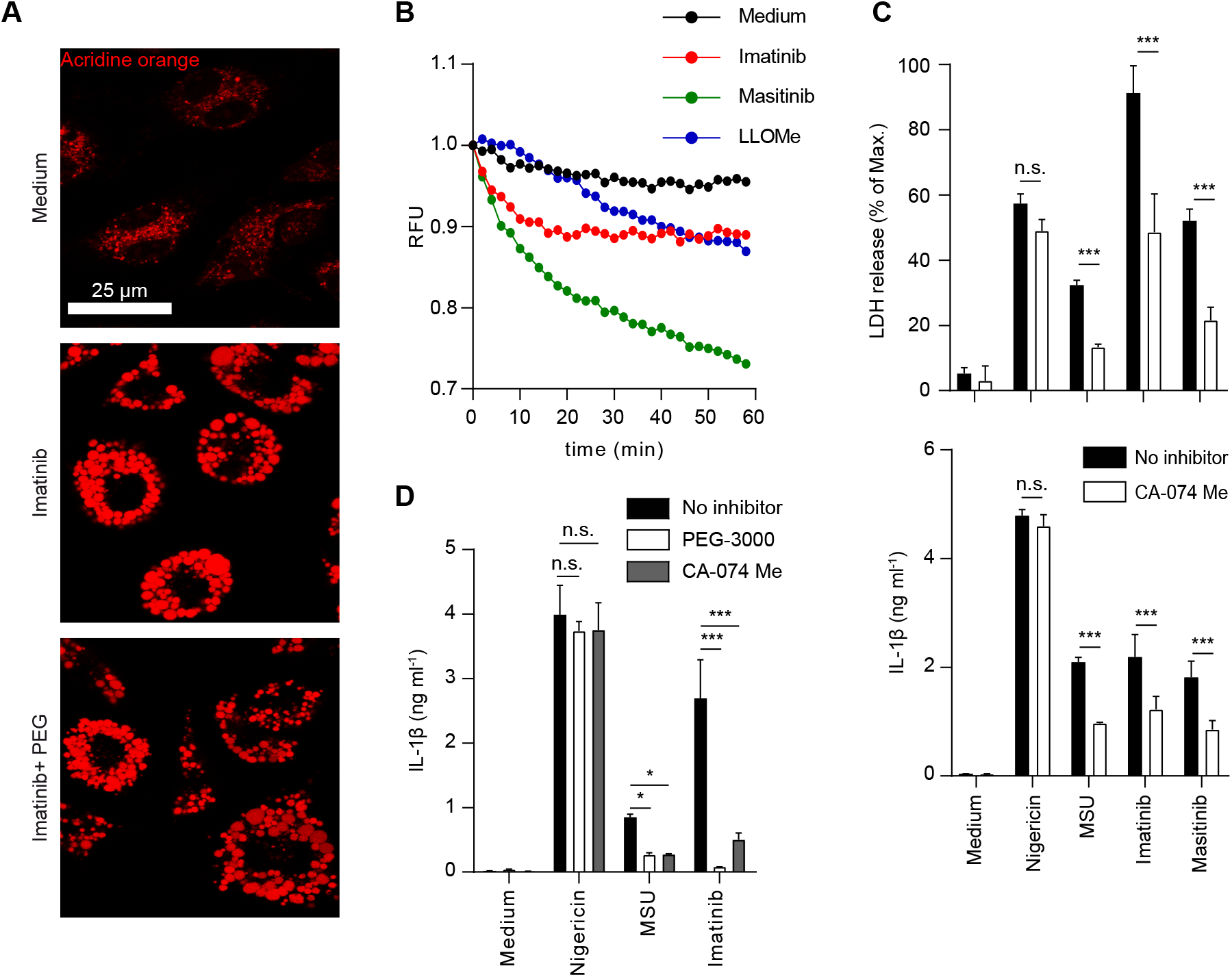
Imatinib and masitinib induce lysosomal damage. (**A**) ASC-deficient *Pycard^-/-^* BMDMs were stained with acridine orange and incubated with 15 mM PEG-3000 or left untreated 30 min prior to stimulation with 40 µM imatinib. Cells were analyzed by fluorescence microscopy after 90 min. (**B**) Acridine orange-stained ASC-deficient *Pycard*^-/-^ BMDMs were stimulated with 40 µM Imatinib, 20 µM Masitinib, 1.25 mM LLOMe or left untreated in the presence of 15 mM PEG- 3000 and subsequently analyzed using a fluorescence plate reader. (**C**) LPS-primed wild-type BMDCs were incubated with 20 µM CA-074Me or left untreated for 1 h prior to stimulation with 5 µM nigericin, 300 µg ml^-1^ MSU, 40 µM imatinib and 20 µM masitinib for 3 h. (**D**) LPS-primed wild-type BMDCs were incubated for 1 h with either 15 mM PEG-3000 or 30 µM CA-074Me, or left untreated. Cells were subsequently stimulated with 5 µM nigericin, 300 µg ml^-1^ MSU or 40 µM imatinib for 3 h. IL-1β secretion and LDH release were determined by ELISA or using a colorimetric assay, respectively, from cell-free supernatants and data are depicted as mean ±SD of technical triplicates. Results are representative of at least three independent experiments. Multiple unpaired T-tests were performed for statistical analysis (*, p < 0.05; **, p < 0.01; ***, p < 0.001; n.s., not significant).

**Fig. 6:**
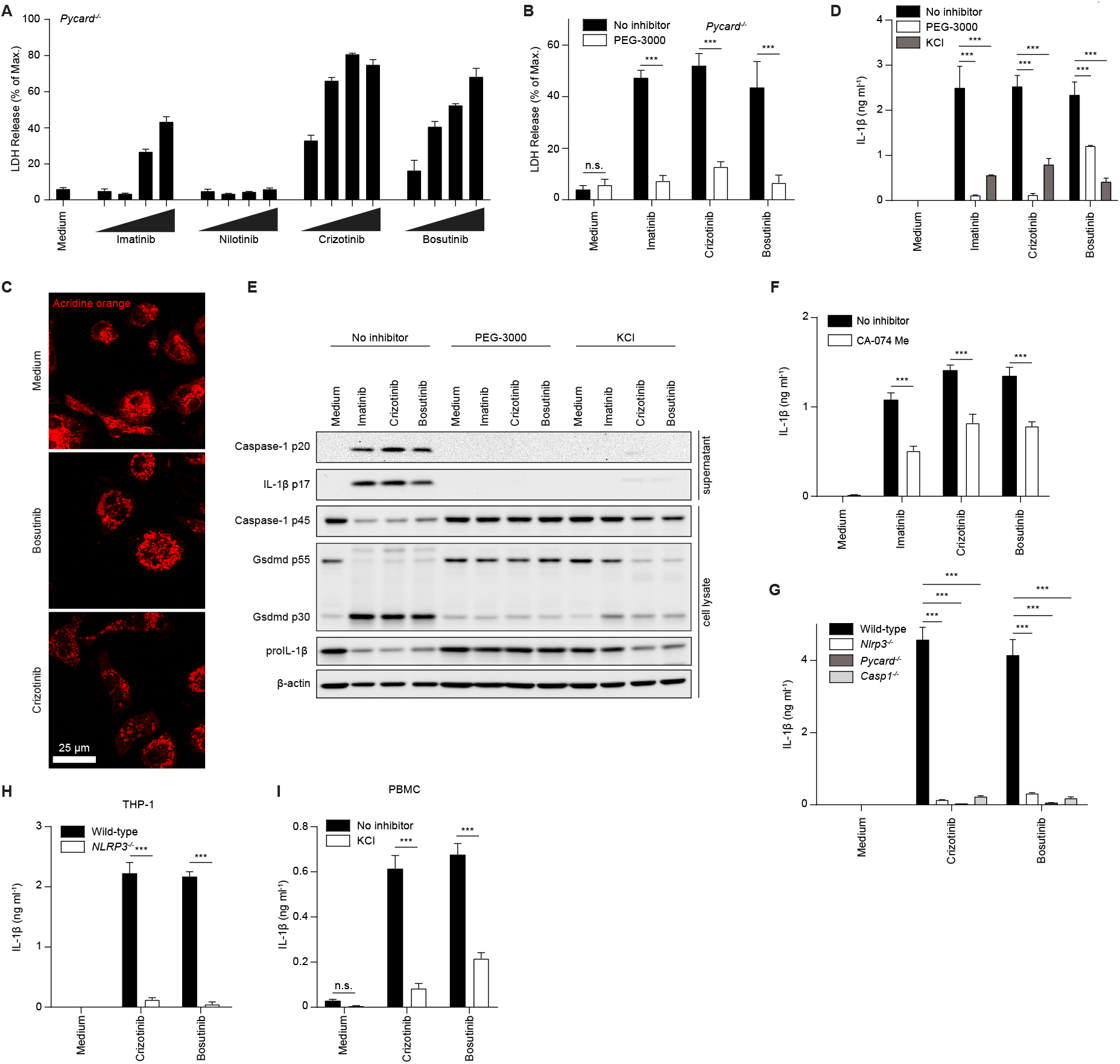
Multiple clinically relevant TKIs induce PEG-sensitive myeloid cell death and NLRP3 inflammasome activation. (**A**) LPS-primed ASC-deficient *Pycard^-/-^* BMDCs were stimulated with 10 µM, 20 µM, 40 µM, and 80 µM of the indicated TKIs for 3 h. (**B**) LPS-primed ASC-deficient *Pycard^-/-^*BMDCs were incubated with 10 mM PEG-3000 or medium 30 min prior to stimulation with 40 µM imatinib, 20 µM crizotinib or 20 µM bosutinib for 3 h. (**C**) ASC-deficient *Pycard^-/-^* BMDMs were stained with acridine orange and incubated with 15 mM PEG-3000 30 min prior to stimulation with 20 µM crizotinib or 40 µM bosutinib. Cells were analyzed by fluorescence microscopy after 10 min. (**D**) and (**E**) LPS-primed wild-type BMDCs were incubated with 10 mM PEG-3000, 50 mM KCl or medium 30 min prior to stimulation with 40 µM imatinib, 20 µM crizotinib or 20 µM bosutinib for 3 h. Cell lysates and supernatants were analyzed by immunoblotting (E). (**F**) LPS-primed wild-type BMDCs were incubated with 30 µM CA-074Me or left untreated for 1 h prior to stimulation with 40 µM imatinib, 20 µM crizotinib or 40 µM bosutinib for 4 h. (**G**) LPS-primed BMDCs from mice of the indicated genotypes were stimulated with 20 µM crizotinib or 20 µM bosutinib. (**H**) THP-1 cells were treated with 200 ng ml^-1^ PMA for 3 h, washed and left at 37 °C, 5 % CO_2_ overnight. Cells were then primed with 50 ng ml^-1^ LPS for 3 h and stimulated with 40 µM crizotinib or 60 µM bosutinib. (**I**) LPS-primed human PBMCs were incubated with 60 mM KCl or medium 30 min prior to stimulation with 10 µM crizotinib or 20 µM bosutinib. IL-1β secretion and LDH release were determined by ELISA or using a colorimetric assay, respectively, from cell-free supernatants and data are depicted as mean ±SD of technical triplicates. Results are representative of at least three independent experiments. Multiple unpaired T-tests were performed for statistical analysis (*, p < 0.05; **, p < 0.01; ***, p <

### Multiple clinically relevant TKIs induce lysosomal swelling, PEG-sensitive myeloid cell lysis, and NLRP3 inflammasome activation

To determine whether other TKIs destabilize myeloid cell membranes and activate NLRP3, we tested several additional molecules of this class (**Supplementary Table 1**) for the induction of lytic myeloid cell death. We found that in addition to imatinib and masitinib, crizotinib (Xalkori^TM^) and bosutinib (Bosulif^TM^) (**Fig. 6A**) as well as fedratinib (Inrebic^TM^), ponatinib (Iclusig^TM^), sunitinib (Sutent^TM^), and dasatinib (Sprycel^TM^) (**Supplementary Fig. 9A**) also induced the rapid release of LDH from ASC- or NLRP3-deficient BMDCs. This cell lysis was inhibited by PEG, suggesting the TKIs share similar mechanisms of cell lysis (**Fig. 6B** and **Supplementary Fig. 9B**). Indeed, like imatinib and masitinib, bosutinib and crizotinib caused lysosomal swelling as visualized by acridine orange (**Fig. 6C**). Consistent with our screening results (where crizotinib killed cells [**Fig. 1B**] and also induced IL-1β release, albeit just below the pre-defined threshold of the screen [**Fig. 1C**]) these TKIs induced substantial release of mature IL-1β from LPS-primed BMDCs (**Fig. 6D** and **Supplementary Fig. 9C**) as well as processing of caspase-1 and GSDMD (**Fig. 6E**). As seen for imatinib and masitinib, PEG, CA- 074Me and extracellular KCl strongly reduced these effects, demonstrating that these TKIs engage the lysosomal pathway for NLRP3 activation (**Fig. 6D-F** and **Supplementary Fig. 9C**). Indeed, IL-1β secretion induced by crizotinib or bosutinib was dependent on NLRP3, ASC and caspase-1, demonstrating that these TKIs are NLRP3 inflammasome activators (**Fig. 6G**). Although several of the TKIs are established inhibitors of the kinase activity of ABL1, ABL2 and the chimeric BCR-ABL1 oncoprotein, there does not seem to be a clear correlation between ABL inhibition in transformed cells and the ability to activate NLRP3 in primary myeloid cells. For example, masitinib, which is primarily a KIT inhibitor and does not inhibit ABL (*48*), does activate NLRP3, while nilotinib is an ABL inhibitor (*49*) but does not cause myeloid cell lysis (**Fig. 1B, 6A**).

To test whether these TKIs also activate human NLRP3, we treated PMA-differentiated THP- 1 cells and LPS-primed primary human PBMCs with the identified TKI inflammasome activators. They also induced substantial IL-1β release from human cells (**Fig. 6H, I** and **Supplementary Fig. 9D),** and this was dependent on NLRP3 (**Fig. 6H** and **Supplementary Fig. 9D**), and inhibited by extracellular KCl as an inhibitor of K^+^ efflux (**Fig. 6I**). Together, these data show that multiple clinically relevant TKIs induce IL-1β release from human myeloid cells by engaging the NLRP3 inflammasome.

## Discussion

Loss-of-function screening strategies aimed at the identification of gene products required for inflammasome activation or at the discovery of small molecule inhibitors have given important insights into how inflammasomes are activated and regulated (*50*). For instance, a genome-wide CRISPR screen identified NEK7 as a factor involved in NLRP3 activation (*50*). However, particularly in the case of NLRP3, screening for activators as a means of interrogating the mechanism of activation might have an important advantage in comparison to loss-of-function approaches. This has to do with the nature of NLRP3, which seems to sense disturbances in cellular homeostasis. If we hypothesize that the mechanism of NLRP3 activation involves sensing disturbances in fundamental cellular processes, then attempts to identify these processes via loss-of-function approaches might fail to report important players because loss of such factors could impair homeostasis or viability in a way that would supersede or conceal any defect in NLRP3 activation as measured by inflammasome-dependent cell death. In contrast, screening for small molecule activators of NLRP3 and identifying their targets is a promising means to reveal upstream mechanisms that might be involved in preserving cellular homeostasis or reporting disturbances to NLRP3.

Here, we screened a library of small molecules and identified new activators of the NLRP3 and NLRP1 inflammasomes. The identification of the antineoplastic drug candidate VbP as an inflammasome activator is consistent with recent studies demonstrating that DPP8/9 inhibitors trigger NLRP1 activation by targeting NLRP1 for partial proteasomal degradation (*25–29*). Indeed, we confirmed that inflammasome activation induced by VbP or the more specific DPP8/9 inhibitor 1G244, which was also found in our screen, was abrogated by the proteasome inhibitor MG132. We believe that expansion of this screening strategy 1) by inclusion of genetic secondary screens (*e.g.* using cells from ASC-deficient mice), 2) by broader exploration of chemical space, and 3) by applying quantitative structure–activity relationship models represent a promising approach for dissecting the mechanism of NLRP3 activation. PEG will also be useful for future screening efforts for NLRP3 inflammasome activators since it can distinguish molecules that activate via the lysosomal pathway from those that trigger K^+^ efflux directly or those that do not require K^+^ efflux, with the latter being of high interest since they might provide long-sought insight into the proximal mechanisms of NLRP3 activation. Precision small molecule activators of NLRP3 (in contrast to currently available activators that either activate NLRP3 by triggering lysis and K^+^ flux or have other pleiotropic effects on myeloid cells), might also have therapeutic value as immunostimulators, for instance in the setting of immunization, infectious disease or cancer immunotherapy, where NLRP3 activation has been shown to have a positive effect for the host (*51*).

A surprising finding of our study is that imatinib, the first molecularly targeted cancer therapy and other TKIs kill and trigger IL-1β secretion from myeloid cells. Imatinib and other TKIs are designed to kill or slow growth of cancer cells addicted to the oncogenes targeted by these small molecules. In BCR-ABL1-positive cancer cells, imatinib can induce caspase-dependent apoptotic death (*52, 53*), but in other cell types it has also been reported to trigger caspase-independent cell death pathways (*54–57*). Here we show that imatinib, masitinib, and other clinically relevant TKIs kill myeloid cells by causing lysosomal swelling and destabilization and vacuolization followed by loss of plasma membrane integrity. The cell death and NLRP3 inflammasome activation mechanism triggered by imatinib and masitinib thus resembles the established lysosomal pathway induced by particulates (such as MSU, alum, silica, cholesterol crystals, and aggregates of endogenous proteins such as β-amyloid or islet amyloid polypeptide) and the lysosomotropic peptide LLOMe (*19, 47*).

An open question is how TKIs trigger lysosomal destabilization. One possibility is that this depends on common ‘off-target’ proteins shared by these TKIs. Examination of the primary kinase targets of the TKIs (*58*) revealed no obvious candidates. Published chemical proteomics studies have uncovered that TKIs simultaneously bind off-target proteins including other kinases and also non-kinases (*59–61*). However, our analysis of a publicly available database (*61*) also did not yield a clear pattern that would explain why some TKIs trigger myeloid cell death and NLRP3 activation while others do not (data not shown). An additional explanation is that physiochemical properties of the TKIs might lead to their accumulation in and disruption of the lysosomal compartment. Indeed, previous studies have shown that certain TKIs including imatinib accumulate in the lysosome (*62, 63*). Such lysosomotropic agents are generally basic and their protonation in the acidic lysosomal lumen decreases their membrane permeability and leads to their accumulation in the lysosome (*64*). In some cases, this can lead to an increase in the osmolarity of the lysosome, and to its swelling and eventual rupture (*65*). The high pkA (*i.e.* weak basic properties) and positive logP (partition coefficient, as an indicator of membrane permeability) of TKIs are consistent with their reported lysosomotropism. Importantly, however, some of the TKIs that do no activate NLRP3 also share these properties. Thus, further work will be required to explain why certain TKIs disrupt the lysosome and activate NLRP3 and others do not.

An intriguing observation is that TKIs triggered cell death specifically in myeloid cells and not in other cell types tested. Myeloid cells are also known as ‘professional phagocytes’ and a high lysosomal capacity is a hallmark of this lineage and a central aspect of their biology. Furthermore, they are sometimes faced with pathogens that escape from or disrupt the lysosome. Mechanisms to sense this disruption and alert neighboring immune cells might mitigate the spread of such pathogens. The myeloid specificity of the cell death triggered by TKIs might be explained by the high lysosomal capacity of these cells, leading to the release of high concentrations of death-inducing factors such as cathepsins. Previous studies suggested the involvement of several lysosomal cathepsins in NLRP3 activation in response to particulates and the lysosomotropic peptide LLOMe (*47, 66*). Inhibition of TKI-induced NLRP3 activation by the cathepsin inhibitor CA-074Me suggests that cathepsins are also involved in NLRP3 activation by these small molecules. The ability of PEG to prevent NLRP3 activation by TKIs (and also by particulates) without blocking lysosomal leakage suggests that lysosomal leakage itself does not directly lead to NLRP3 activation. However, the ability of PEG to block cell lysis and NLRP3 activation in response to TKIs, as well as the requirement for K^+^ efflux suggest that TKI-induced lysosomal disruption causes NLRP3 activation by inducing plasma membrane damage and K^+^ efflux. Notably, PEG also blocks cell lysis and NLRP3 activation induced by particulates such as MSU, again pointing to the central role of lytic K^+^ efflux in NLRP3 activation via the lysosomal pathway. Since lysosome-disrupting particulates are implicated as key activators of NLRP3 in inflammatory disease, this finding provide insight into the mechanism of NLRP3 activation in disease. PEG also provides an experimental tool to uncouple lysosomal swelling from cell death and therefore to investigate the mechanism by which particulates and lysosomotropic molecules trigger cell death and NLRP3 activation.

The finding that imatinib and masitinib as well as other TKIs can kill myeloid cells and cause release of pro-inflammatory mediators raises the question as to whether this might contribute to their efficacy or adverse effects. The rapid lytic cell death we observed in primary myeloid cells in response to imatinib and masitinib did not occur in BCR-ABL1–positive cell lines, suggesting it does not directly contribute to elimination of cancer cells. However, the ability of selected TKIs to kill myeloid cells could potentially contribute to reported effects of TKIs such as myelosuppression, neutropenia, inhibition of dendritopoiesis, reduction of tumor-associated M1 macrophages, diarrhea, or increased susceptibility to infection (*67, 68*). Among TKIs, imatinib in particular has been reported to have pleiotropic immunological effects (*69*), to which the IL-1β we observed might contribute. IL-1β can display pro- or anti-neoplastic effects (*67–69*), but in hematological malignancies (*70*) and especially in chronic myelogenous leukemia (which is a major therapeutic setting for imatinib) appears to promote disease progression (*71, 72*). Therefore, the NLRP3-dependent cytokine secretion induced by imatinib probably does not significantly contribute to its clinical efficacy. A major consideration in evaluating whether our findings are relevant in the therapeutic setting is that the minimum doses of TKIs required to trigger lytic cell death and inflammasome activation exceed those generally needed to inhibit the kinase. Imatinib inhibits BCR-ABL1 activity with a IC_50_ ranging from 0.12- 0.47 µM (*73, 74*) *in vitro* and complete inhibition is achieved at 1 µM (*52, 53*). Though pharmacokinetic studies suggest that the mean steady-state plasma concentration for a daily imatinib dose of 400 mg is about 2 µM, imatinib reaches average peak plasma concentrations of 5 µM, with higher peak concentrations in the plasma of individual patients (*75, 76*). Although 5 µM is somewhat below the range at which TKIs induce cell death and IL- 1β secretion in cell culture, it is possible that local peak concentrations (*e.g.* in the upper gastrointestinal tract or liver for orally administered TKIs) may be sufficient to trigger membrane destabilization or NLRP3 activation in resident myeloid cells or potentially other cell types. However, it is clear that further studies will be needed to determine whether TKIs trigger lytic cell death and NLRP3 activation in the therapeutic setting, and if so, whether this influences patient outcomes.

## Materials and Methods

### Mice

*Nlrp3^-/-^* (*30*), *Pycard^-/-^*(*77*), *ICE^-/-^* (*Casp1/11^-/-^*) (*78*), *Casp1^-/-^* (*32*), *Casp1^mlt/mlt^* (*32*), *Gsdmd^-/-^* (*32*), ASC^citrine^ (*79*), and wild-type mice of C57BL/6 and BALB/c backgrounds were housed under SOPF or SPF conditions at the Center for Experimental Models and Transgenic Services (CEMT, Freiburg, Germany), the Zentrum für Präklinische Forschung (ZPF, Munich, Germany), Charles River Laboratories (Calco, Italy), or the Center of Infection and Immunity (University of Lausanne, Epalinges, Switzerland) in accordance with local guidelines.

### Cell lines

Cell lines were cultured in T75 flasks at 37°C, 5% CO_2_ in a humidified incubator and continuously passaged. Media were supplemented with 10% fetal calf serum (FCS) (Gibco) and 100 U ml^-1^ penicillin-streptomycin (Gibco). HeLa, HEK 293T, NIH-3T3 and mouse embryonic fibroblasts (MEF) were cultured in DMEM (Gibco) and MEF medium was additionally supplemented with 0.1% β-mercaptoethanol (Gibco). THP-1 cells were cultured in RPMI (Gibco) and HCT 116 cells were grown in McCoy’s 5a Medium (Gibco).

### BMDC and BMDM Preparation and Stimulation

Cells were cultured at 37°C, 5% CO_2_ in a humidified incubator. Murine bone marrow-derived dendritic cells (BMDCs) and macrophages (BMDMs) were differentiated from tibial and femoral bone marrow as previously described in detail (*80*). Recombinant human M-CSF and murine GM-CSF (Immunotools) were used at 100 ng ml^-1^ and 20 ng ml^-1^, respectively. After 6 - 8 days of differentiation, cells were plated in 96-well plates at a density of 0.8 - 1.5×10^5^ cells per well, primed with 20 - 150 ng ml^-1^ *E. coli* K12 ultra-pure LPS (InvivoGen) for 2 - 3 h, and treated with inflammasome activators, TKIs and other stimuli for 0.5 - 16 h. All stimulations were performed in triplicates and cytokine production in cell-free supernatants was measured by ELISA. TKIs were purchased from Sellekchem and treatment was typically performed at 20 µM, 40 µM, 60 µM, and 80 µM. Other inflammasome activators and stimuli were used as follows: 5 mM ATP (Sigma), 5 μM nigericin (Sigma), 300 μg ml^-1^ MSU (prepared as previously described (*30*)), 1 - 2 μg ml^-1^ poly(dA:dT) (InvivoGen) (transfected with Lipofectamine 2000, Invitrogen), 100 µM imiquimod (R837) (InvivoGen), 100 µM CL097 (InvivoGen), 1 - 10 µM VbP (MedChem Express), 1 - 10 µM 1G244 (Sigma), 1 - 10 µM alogliptin (MedChem Express), 10 µM raptinal (Adipogen), 1 µM (1S,3R)-RSL3, and 1 mM H_2_O_2_ (PHC Corporation). Inhibitors were added after 2 - 2.5 h of priming, and 20 - 60 min before stimulation with inflammasome activators. Inhibitor concentrations were typically used at the lowest dose showing robust, reproducible efficacy: 20 - 40 μM zVAD-fmk (Enzo), 20 - 60 mM KCl (Sigma), 3 - 5 μM MCC950 (Adipogen), 50 - 200 nM MG132 (Sigma), 3 µM cytochalasin D (Sigma), 50 - 150 mM PEG-600, 5 - 15 mM PEG-3000 (Sigma and Merck), 20 - 30 µM CA-074Me (Calbiochem), 0,4 µM ferrostatin-1 (Biomol), 10 µM PJ-34 (MedChemExpress) 50 mM glycine (Labochem) and 50 mM sucrose (Sigma). To minimize off-target effects of extracellular KCl, it was added and mixed well by pipetting immediately before addition of inflammasome activators.

### PBMC isolation

Peripheral blood samples were diluted 1:1 with PBS containing 2 mM EDTA, and human Pancoll, density 1077 g ml^-1^ (Pan Biotech) was layered underneath. It was centrifuged for 30 min at 475 x g, 21°C, the upper plasma layer was removed and the mononuclear cell layer carefully transferred to PBS-EDTA. After 15 min of centrifugation, the supernatant was completely removed and the cell pellet resuspended in red blood cell lysis buffer (BioLegend). The cells were washed and then cultured for at least 2 h in RPMI with 10% FCS, 100 U ml^-1^ penicillin-streptomycin. Typically, 0.25×10^6^ cells per well were plated in 96-well plates. The cells were then again washed and finally cultured in RPMI with 10% FCS, 100 U ml^-1^ penicillin-streptomycin and 100 ng ml^-1^ recombinant human M-CSF. After cultivation for at least 24 h, the cells were used for inflammasome stimulation (see *BMDC and BMDM Preparation and Stimulation*).

### Primary thymocyte isolation

Primary murine thymocytes were isolated by removing the thymus from euthanized mice and mashing it through a 100 µm nylon cell strainer. Single cell suspensions were collected in RPMI with 10% FCS, 100 U ml^-1^ penicillin-streptomycin and centrifuged for 5 min at 400 x g, 4°C. Erythrocytes were lysed using red blood cell lysis buffer (Biolegend) and the reaction was stopped by addition of medium. For stimulation with TKIs and LDH release assay, 0.8×10^6^ cells per well were plated to 96-well plates.

### Screening for inflammasome activators

To identify small molecule NLRP3 activators, a 2-step high-throughput screening of a public Novartis small molecule library (*33*) was conducted. For primary screening, 200 nl of the small molecule stock solution (10 mM in DMSO) were transferred to white 384-well microplates (Greiner) using an Echo 555 Acoustic Technology Liquid Handler (Labcyte). BMDCs on day 7 of differentiation were primed with 50 ng ml^-1^ LPS and immediately plated in 40 µl medium to the small molecule-containing 384-well plates with a Multidrop Combi Reagent Dispenser (Thermo Fisher Scientific) at 0.25×10^6^ cells ml^-1^. The final small molecule concentration was 50 µM. Following 20 h of incubation at 37°C, 5% CO_2_, reduction in cell viability was determined as reduction of cellular ATP levels by using the CellTiter-Glo Luminescent Cell Viability Assay (Promega). Cells were treated with 5 µM nigericin as a positive control for inflammasome activation or 0.5% DMSO as a negative control. Compound testing was performed as single point measurements, whereas 8 wells of positive or negative control each were included per plate. To compare cell viability, the lowest luminescence signal obtained was subtracted from all signals and all luminescence signals were then calculated as % of the highest luminescence signal. Small molecules that reduced cellular ATP levels to less than 50% were considered active and therefore included in the functional secondary screen for IL-1β release. For that, BMDCs on day 7 were treated with 50 ng ml^-1^ LPS, directly plated to white 384-well microplates and primed for 3 h at 37°C, 5% CO_2_. Subsequently, using an Echo 555 Acoustic Technology Liquid Handler, 200 nl of the small molecule stock solutions (10 mM in DMSO) were transferred to the microplates giving a final concentration of 50 µM and it was incubated for 20 h at 37°C, 5% CO_2_. Cell viability was assessed as in the primary screen and IL-1β in the cell- free supernatants was detected using a commercial mouse IL-1β homogenous time-resolved fluorescence (HTRF) kit (Cisbio) according to the manufacturer’s instructions. In brief, anti-IL-1β cryptate antibody and anti-IL-1β d2 antibody were prediluted in detection buffer and mixed in equal parts. 10 µl antibody mix was transferred to black, small volume 384-well microplates (Greiner) with a Microlab STAR Liquid Handling System (Hamilton), 10 µl cell-free supernatants were added and it was incubated overnight at 4°C. Fluorescence signals were measured in a time-resolved manner. Again, cells were treated with 5 µM nigericin as a positive control or 0.5 % DMSO as a negative control. HTRF signals for 0.5% DMSO were subtracted from all values and 5 µM nigericin was set to 100% to assess IL-1β signal intensity.

### Immunodetection of Proteins

For cytokine quantification of cell-free supernatants, ELISA kits for murine and human IL-1β, IL-1α and TNF (eBioscience or Invitrogen) were used. ELISA data is depicted as mean ±SD of technical triplicates as previously described (*80*). For immunoblot analysis, cell-free supernatant and cell lysate samples in SDS- and DTT-containing sample buffer were analyzed. Triplicates were pooled and proteins were separated by SDS-PAGE and transferred to nitrocellulose membranes using standard techniques (*32*). The following primary antibodies were used: anti-Caspase-1 (p20) mAb (Casper-1, Adipogen), IL-1β/IL-1F2 pAb (R&D Systems), anti-GSDMD (EPR19828, Abcam), anti-NLRP3/NALP3 mAb (Cryo-2, Adipogen), anti-Asc pAb (AL177, Adipogen), β-Actin mAb (8H10D10, #3700, Cell Signaling Technology), anti-HMGB1 antibody (ab18256, Abcam), and anti-α-tubulin mAb (T5168, Sigma).

### Cell Viability Assays

Lytic cell death was determined by measuring LDH release from cell-free supernatants using a colorimetric assay (Promega or Takara) according to the manufacturer’s protocol. Medium served as blank value and was subtracted from the sample values. Results were plotted as percentage of 100% dead cells lysed with lysis buffer 45 min prior to collection of the cell supernatants. Total cellular ATP was measured using a luminescent assay (CellTiter-Glo, Promega) according to the manufacturer’s instructions. Data is depicted as mean ±SD of technical triplicates

### Acridine orange staining of lysosomes

BMDMs were plated at a density of 0.1×10^6^ cells per well in their regular medium into black-walled 96-well plates and stained for 15 min with acridine orange (Invitrogen) according to manufacturers instructions. The acridine orange containing medium was removed, cells were washed once with assay buffer consisting of PBS with 2.5% FCS, and assay buffer additionally containing 15 mM PEG-3000 was added. Cells were stimulated as indicated and fluorescence at λ_ex_ 475- 495 nm and λ_em_ 525- 545 nm was recorded every 2 minutes. Data is depicted as mean of technical triplicates.

### Fluorescence Imaging

BMDMs were plated at a density of 0.05 - 0.1×10^6^ cells per well in 8-well µ-slides (IbiTreat, Ibidi). Cells were primed with 50 ng ml^-1^ LPS for 2 h followed by treatment with compounds as indicated. Cells were washed with PBS, fixed in 4% paraformaldehyde (PFA) for 10 min and permeabilized in PBS with 0.1% (v/v) Triton X-100 for 5 min. Cells were stained with anti-ASC primary antibody (AL177, Adipogen) diluted in blocking buffer consisting of PBS, 5% FCS and 0.1% Triton X-100, followed by anti-rabbit IgG cross-absorbed secondary antibody (Alexa Fluor 555, Invitrogen), and finally mounted in Vectashield antifade mounting medium containing DAPI (Vector Laboratories).

For live cell imaging of unfixed cells at 37°C in a 5% CO_2_ humidified atmosphere, the cells were primed with 50 ng ml^-1^ LPS for 2 h or left unprimed and then stained as indicated. The following stains were used: Vybrant Dil cell-labeling solution (Invitrogen), Draq7 (BioLegend), Annexin V-FITC apoptosis staining/detection kit (Abcam, BioLegend) and acridine orange (Invitrogen). Hoechst 3342 (Invitrogen) was used to stain the nuclei.

Confocal microscopy was performed with a Leica SP8 confocal microscope equipped with a 63×/ 1.40 and 40×/ 1.25 oil objective (Leica Microsystems) keeping the laser settings of the images constant for each experiment to allow direct comparison of signal intensities between images of the same channel.

### K^+^ Measurement

Intracellular K^+^ measurements were performed by reflection X-ray fluorescence analysis (TXRF) as described previously (*16*). Cells were stimulated in 96-well plates. After supernatants were removed, the residual medium was completely aspirated. The cells were extracted by adding 25 μl of an ultra-pure 3 % dilution of HNO_3_ in water containing 5 μg ml^-1^ vanadium as internal standard to the wells. 5 μl lysates were spotted on a silicon wafer and evaporated to dryness. Measurement was performed with an Atomika TXRF 8010 device equipped with a molybdenum x-ray tube. Characteristic signals for potassium (EKα = 3,31 keV) and vanadium (EKα = 4,95 keV) were used for data evaluation using the software Spectra Picofox (Bruker).

### Analysis of total phospho-Tyrosine

K-562 cells were plated to non-tissue culture treated 96-well plates and treated with PEG and TKIs. After incubation, cells were spun down, resuspended in PBS and transferred to 96-well V-bottom plates. Cells were washed with PBS and stained with Zombie Aqua fixable viability dye (BioLegend) according to the manufacturer’s instructions. Cells were then washed once with FACS buffer (PBS with 2% FCS) and fixed with 2% PFA. PFA was removed by centrifugation and the cells resuspended in PBS. To permeabilize the cells, 100% methanol was slowly added to the pre-chilled cells on ice to a final concentration of 90% methanol before a 30 min incubation on ice. Finally, intracellular immunostaining for total phospho-Tyrosine was performed by incubating with phospho-tyrosine mAb (P-Tyr-100 Alexa Fluor 647, Cell Signaling Technology). Cells were washed twice with and resuspended in FACS buffer and analyzed using a BD FACS Canto II flow cytometer (BD Biosciences). Data were acquired with DIVA (BD Biosciences) and analyzed with FlowJo software (FlowJo LLC, BD).

### Cell death characterization by flow cytometry

Pacific Blue Annexin V Apoptosis Detection Kit with 7-AAD (BioLegend) was used to characterize cell death by flow cytometry. To this end, BMDCs were treated with TKIs as indicated, harvested with HBSS/EDTA, and transferred to 96-well V-bottom plates. Cells were stained with Pacific Blue Annexin V and 7-AAD in Annexin V binding buffer according to the manufacturer’s protocol. The cells were washed and analyzed with a BD FACS Canto II (BD Biosciences) flow cytometer. Data were acquired with DIVA (BD Biosciences) and were analyzed with FlowJo software (FlowJo LLC, BD).

### Transmission electron microscopy

After 7 days of cultivation, 10^7^ BMDMs in 10 ml medium were transferred to 50 ml conical centrifugation tubes and incubated with 40 µM imatinib, 20 µM masitinib or 0.05% DMSO for 2 h at 37°C, 5% CO_2_. During incubation, the tubes were inverted every 15 min to prevent attachment of the cells to the walls of the tube, and to ensure equal distribution of the added compounds. Glutaraldehyde (GA) was added to a final concentration of 1% and the cells were fixed for a maximum of 10 min at 37°C, 5% CO_2_. The cell suspension was centrifuged for 5 min at 400 x g, 4°C and the cell pellet was resuspended in HEPES buffer with 1% GA and further fixed for 2 h at room temperature. The fixed cells were then left in suspension at 4°C overnight.

Cells were collected by centrifugation at 400 x g for 5 min and cell pellets were embedded in 2% low melting agarose. Pellets were washed five times for 10 min each with HEPES buffer at room temperature and post fixed for 2 h at 4°C with a 1% aqueous solution of OsO_4_. Cells were five times washed with distilled H_2_O for 10 min each and *en bloc* stained with 1% uranyl acetate in water for 1 h at room temperature. Dehydration in increasing graded series of ethanol from 30% to 95% (10 min per change) followed by 100% ethanol twice, and 100% acetone twice for 30 min was carried out before embedding in Epon 812 resin. Sections of 70 nm were obtained using a Reichert-Jung ultramicrotome and collected in slot grids. After post staining with 2% uranyl acetate and Reynolds lead-citrate stain, sections were observed in a Philips CM-10 (80 kV) electron microscope equipped with a Gatan Bioscan 792 camera or with a Hitachi TEM 7800 (80 kV) equipped with an EMSIS Xarosa camera.

### Global analysis of hits as compared to the whole library

Novartis small molecule library annotation data (manually adjusted for library composition used for screening (*33*)) was extended using the webchem R package (DOI:10.18637/jss.v093.i13 (1.1.0 with R 3.6.3)) and manual curation.

## Acknowledgements

The authors thank the animal caretakers at University Medical Center Freiburg and Klinikum rechts der Isar Munich for their support, and Susanne Weiß, Ina Spirer, Valentin Höfl, Gerrit Siegers, Caroline Schwenzel, Nico Wanjura, Timo Kleindienst, Shaumya Kulendran, Rosula Hinnenberg, and Natacha Stoehr for technical assistance. Klaus-Peter Knobeloch provided primary MEFs, Romeo Ricci provided NLRP3-deficient THP-1 cells, Lena Illert provided K- 562 cells, Petr Broz provided *Gsdme^-/-^* BMDMs. We also thank Robert Huber, Ruth Geiss-Friedlander, Guillaume Médard, Robert Zeiser, Marco Prinz, Angelika Rambold, Anne-Kathrin Classen and Paul Manley for helpful discussions.

## Funding

This work was supported by the Deutsche Forschungsgemeinschaft (DFG, German Research Foundation) through SFB 1160 (Project ID 256073931), SFB/TRR 167 (Project ID 259373024), SFB 1425 (Project ID 422681845), SFB 1479 (Project ID 441891347) (to O.Gr.), SFB 850 Project B7 (to T.R.), GRK 2606 (Project ID 423813989) (to O.Gr. and T.R.), SFB 1335 (Project ID 360372040) (to P.J.J.), and under the Germany’s Excellence Strategy (CIBSS - EXC-2189 - Project ID 390939984, to O.Gr., R.B. and T.O.), as well as by the European Research Council (ERC) through Starting Grant 337689 and Proof-of-Concept Grant 966687 (to O.Gr.), and the German Consortium for Translational Cancer Research (DKTK) (to T.R.). The TEM (Hitachi HT7800) was funded by the DFG grant INST 39/1153-1 and is operated by the University of Freiburg, Faculty of Biology, as a partner unit within the Microscopy and Image Analysis Platform (MIAP) and the Life Imaging Center (LIC), Freiburg.

## Author Contributions

E.N., G.M., T.C., A.K., S.W., B.S.S., N.F., O.Go., and C.J.G. performed experiments and analyzed data, prepared figures, and wrote figure legends and the methods section of the manuscript. C.A., C.P., A.B., and C.J.F. helped design, perform and analyze the screen. M. R.-F. generated EM data. S.F., S.R. and R.B. performed bioinformatics analysis. T.O., M.T. and T.R. helped design and interpret a portion of the experiments. O.Go. and P.J.J. supervised a portion of the work. O.Gr. and C.J.G. wrote the main text of the manuscript, with the help of E.N. and O.Go. O.Gr. conceived and oversaw the project.

## Competing interests

S.R., C.A., C.P., A.B., and C.J.F. are employees of Novartis, Basel, Switzerland. P.J.J. has had a consulting or advisory role, received honoraria, research funding, and/or travel/accommodation expenses not related to the present work from: Ariad, Abbvie, Bayer, Boehringer, Novartis, Pfizer, Servier, Roche, BMS and Celgene, Pierre Fabre, Janssen / Johnson&Johnson, MSD.

