## Supplemental Figures 1-9, Supplemental Table 1 for "Tyrosine kinase inhibitors trigger lysosomal damage-associated cell lysis to activate the NLRP3 inflammasome"

Neuwirt et. al.

### Supplementary Material

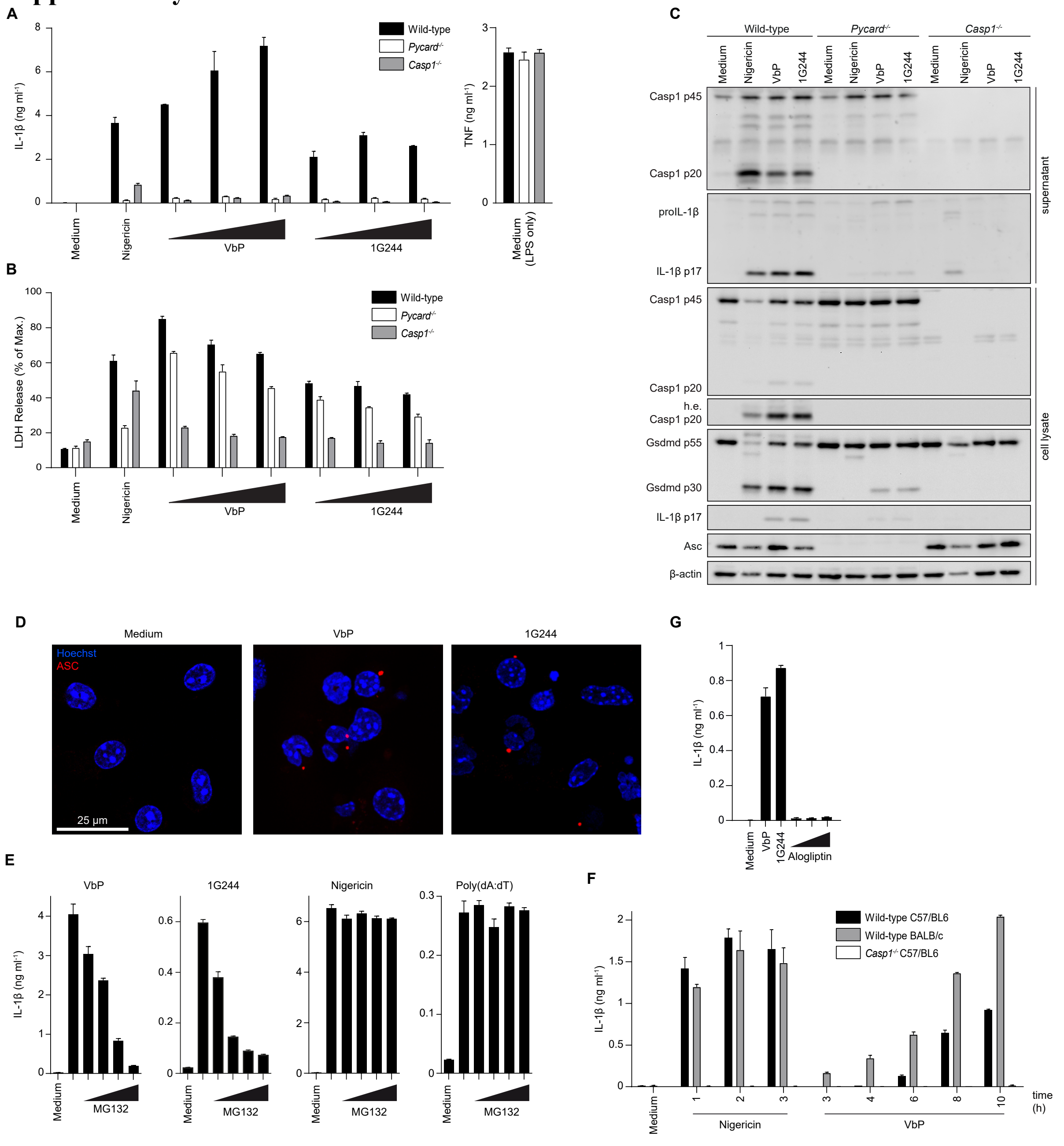


**Supplementary Fig. 1: Inflammasome activation by dipeptidyl peptidase inhibitors**

(**A**), (**B**) and (**C**) LPS-primed wild-type, ASC (*Pycard*$^{-/-}$) and Caspase-1-deficient BMDCs were stimulated with 5 µM nigericin or increasing concentrations of VbP (1, 3, 10 µM) and 1G244 (1, 3, 10 µM) for 10 h. IL-1β and TNF secretion (A) and LDH release (B) were determined and cell lysates and supernatants were analyzed by immunoblotting for the given proteins (10 µM shown for VbP and 1G244) (C).

(**D**) Confocal microscopy of LPS-primed *Casp1*$^{mlt/mlt}$ BMDMs stimulated with 10 µM VbP or 3 µM 1G244 for 8 h. Nuclei were localized with Hoechst 33342 (blue) and ASC specks stained with an anti-ASC antibody (red).

(**E**) LPS-primed wild-type BMDCs were incubated with MG132 (50, 100, 150 and 200 nM) 30 min prior to stimulation with 10 µM VbP, 3 µM 1G244, 5 µM nigericin or 1 µg ml$^{-1}$ poly(dA:dT) for 8 h and IL-1β secretion was measured.

(**F**) LPS-primed BMDCs from wild-type and *Casp1*$^{-/-}$ C57/BL6 mice or wild-type BALB/c mice were stimulated with 5 µm nigericin or 10 µM VbP for the indicated times and IL-1β secretion was measured.

(**G**) LPS-primed wild-type BMDCs were stimulated with 10 µM VbP, 3 µM 1G244 or alogliptin (1, 3, 10 µM) for 10 h.

Cytokine secretion and LDH release were determined by ELISA or using a colorimetric assay, respectively, from cell-free supernatants and data are depicted as mean ±SD of technical triplicates. Results are representative of at least three independent experiments.

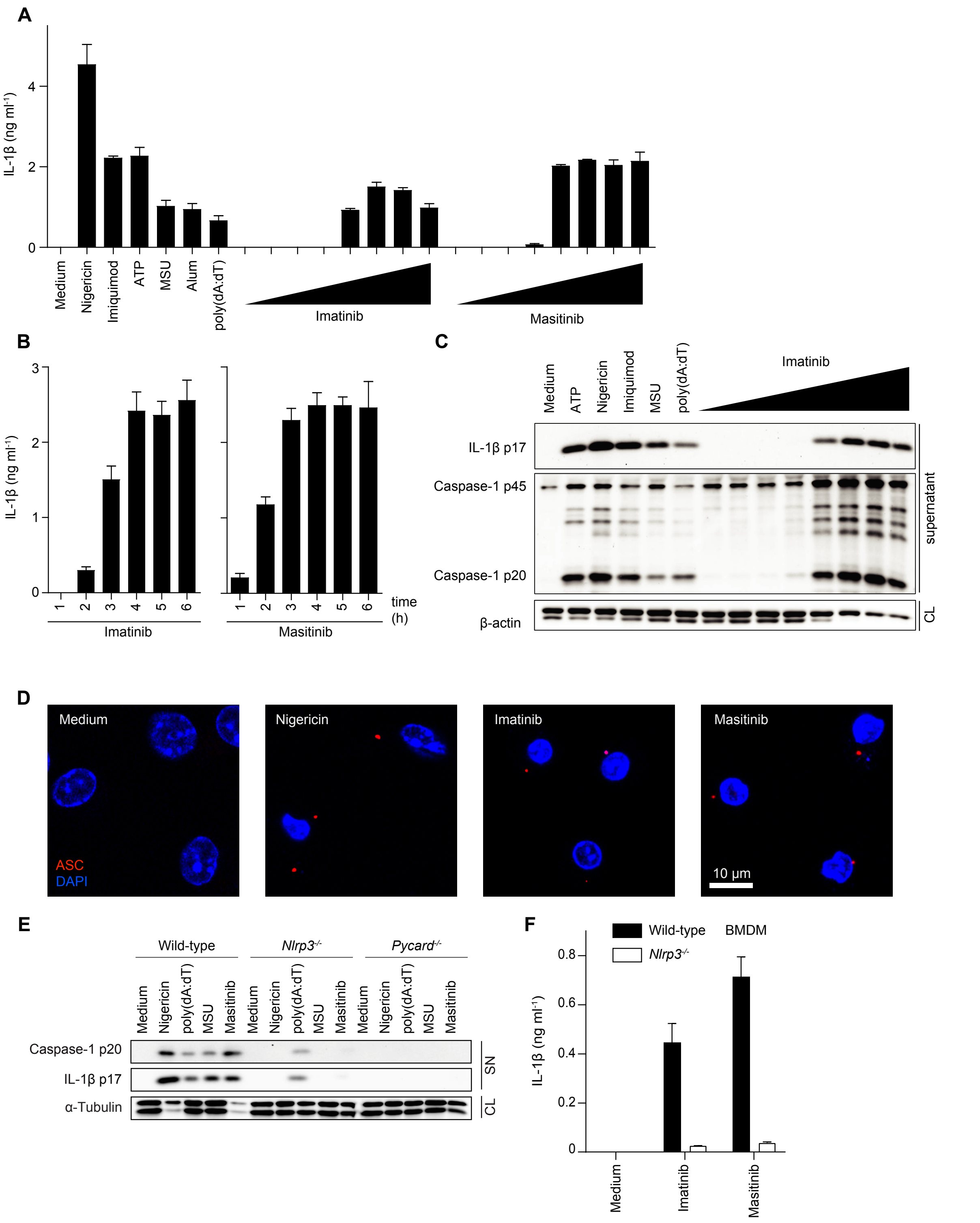


**Supplementary Fig. 2: Supplementary data corresponding to main Fig. 2: Imatinib and masitinib activate the NLRP3 inflammasome**

(**A**) LPS-primed wild-type BMDCs were stimulated with imatinib and masitinib (1, 5, 10, 20, 40, 60, 80, 100 µM) for 3 h and 5 µM nigericin, 100 µM imiquimod, 5 mM ATP, 300 µg ml$^{-1}$ MSU or alum and 2 µg ml$^{-1}$ poly(dA:dT) and IL-1β secretion was measured.

(**B**) IL-1β secretion time course from LPS-primed BMDCs treated with 60 µM imatinib and masitinib and IL-1β secretion was measured.

(**C**) Immunoblot analysis of cell lysates and cell-free supernatants from (A).

(**D**) Confocal microscopy of LPS-primed BMDCs stimulated with 5 µM nigericin, 60 µM imatinib or 40 µM masitinib. Nuclei were localized with DAPI (blue) and ASC specks stained with an anti-ASC antibody (red).

(**E**) Immunoblot analysis for Caspase-1 and IL-1β in cell-free supernatants from LPS-primed wild-type, *Nlrp3*$^{-/-}$ or ASC-deficient *Pycard*$^{-/-}$ BMDCs stimulated with inflammasome activators as indicated. Cell lysates were analysed for tubulin.

(**F**) LPS-primed BMDMs from wild-type and NLRP3-deficient mice were incubated with 60 mM KCl 30 min prior to stimulation with 40 µM imatinib or 20 µM masitinib for 3 h and IL-1β secretion was measured.

IL-1β secretion was determined by ELISA from cell-free supernatants and data are depicted as mean ±SD of technical triplicates. Results are representative of at least three independent experiments.

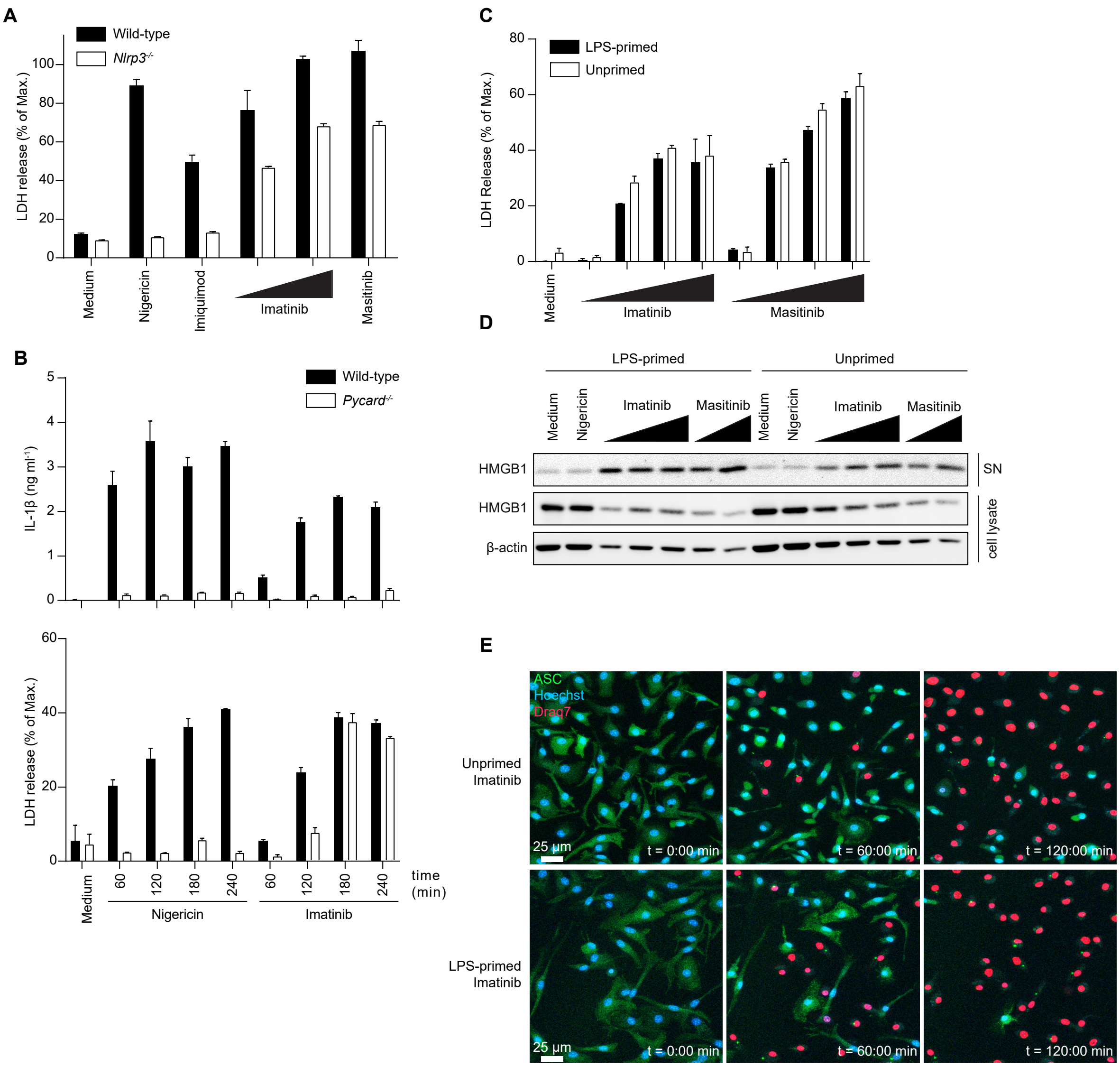


**Supplementary Fig. 3: Supplementary data corresponding to main Fig. 2: Imatinib and masitinib trigger inflammasome-independent lytic cell death**

(**A**) LPS-primed wild-type and NLRP3-deficient BMDCs were stimulated with 5 µM nigericin, 100 µM imiquimod, 40 and 60 µM imatinib or 40 µM masitinib and LDH release was measured.

(**B**) LPS-primed wild-type and ASC-deficient *Pycard*$^{-/-}$ BMDCs were stimulated with 5 µM nigericin or 40 µM imatinib as indicated and IL-1β secretion and LDH release were measured.

(**C**) BMDCs from ASC-deficient *Pycard*$^{-/-}$ mice were primed with 50 ng ml$^{-1}$ LPS for 3 h or left unprimed. Cells were then stimulated with increasing concentrations of imatinib and masitinib (20, 40, 60, 80 µM) for 3 h and LDH release was measured.

(**D**) Immunoblot analysis of cell lysates and supernatants from (C) (5 µM nigericin, 40, 60, 80 µM imatinib, 40, 60 µM masitinib) for HMGB1 release.

(**E**) BMDMs from ASC$^{citrine}$ reporter mice were left unprimed or primed with 50 ng ml$^{-1}$ LPS for 2 h. Cells were stained with Hoechst 33342 (blue) and DRAQ7 (red), stimulated with 40 µM imatinib and analyzed by fluorescence microscopy after 0, 60 and 120 min.

IL-1β secretion and LDH release were determined by ELISA or using a colorimetric assay, respectively, from cell-free supernatants and data are depicted as mean ±SD of technical triplicates. Results are representative of at least three independent experiments.

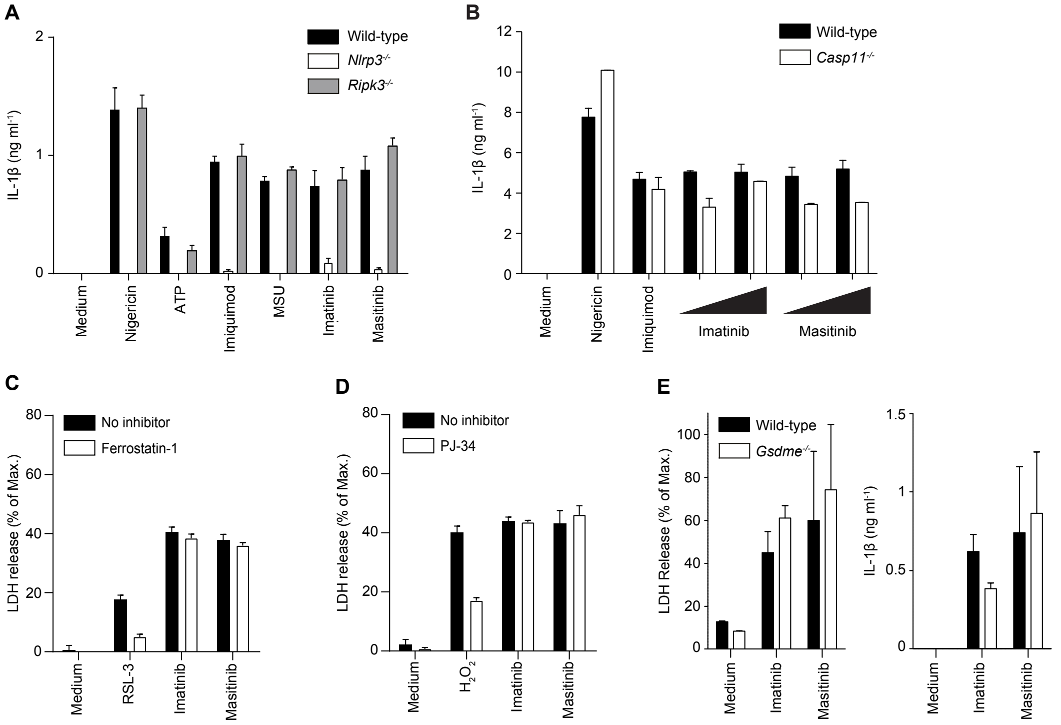


**Supplementary Fig. 4: TKI-induced lytic cell death is not necroptosis, pyroptosis, ferroptosis, or parthanatos**

(**A**) LPS-primed BMDCs from wild-type, *Nlrp3*$^{-/-}$ and *Ripk3*$^{-/-}$ mice were stimulated with imatinib (40 µM), masitinib (20 µM) and control activators as indicated for 3 h and IL-1β secretion was measured.

(**B**) LPS-primed BMDCs from wild-type or caspase-11-deficient mice were stimulated with 5 µM nigericin, 100 µM imiquimod, imatinib (40, 60 µM) or masitinib (20, 40 µM) and IL-1β secretion was measured.

(**C**) and (**D**) ASC-deficient *Pycard*$^{-/-}$ BMDCs were incubated with 0,4 µM Fer-1 (C) or 10 µM PJ-34 (D) 1 h prior to stimulation with 1 µM RSL-3 (C) or 1 mM $H_2O_2$ (D) and 60 µM imatinib or 40 µM masitinib for 5 h and LDH release was measured. Cells were left unprimed since TLR stimulation blocks ferroptosis and parthanatos.

(**E**) LPS-primed wild-type or *Gsdme*$^{-/-}$ BMDMs were stimulated with 40 µM imatinib or 20 µM masitinib for 4 h and IL-1β secretion and LDH release were measured.

IL-1β secretion and LDH release were determined by ELISA or using a colorimetric assay, respectively, from cell-free supernatants and data are depicted as mean ±SD of technical triplicates. Results are representative of at least three independent experiments.

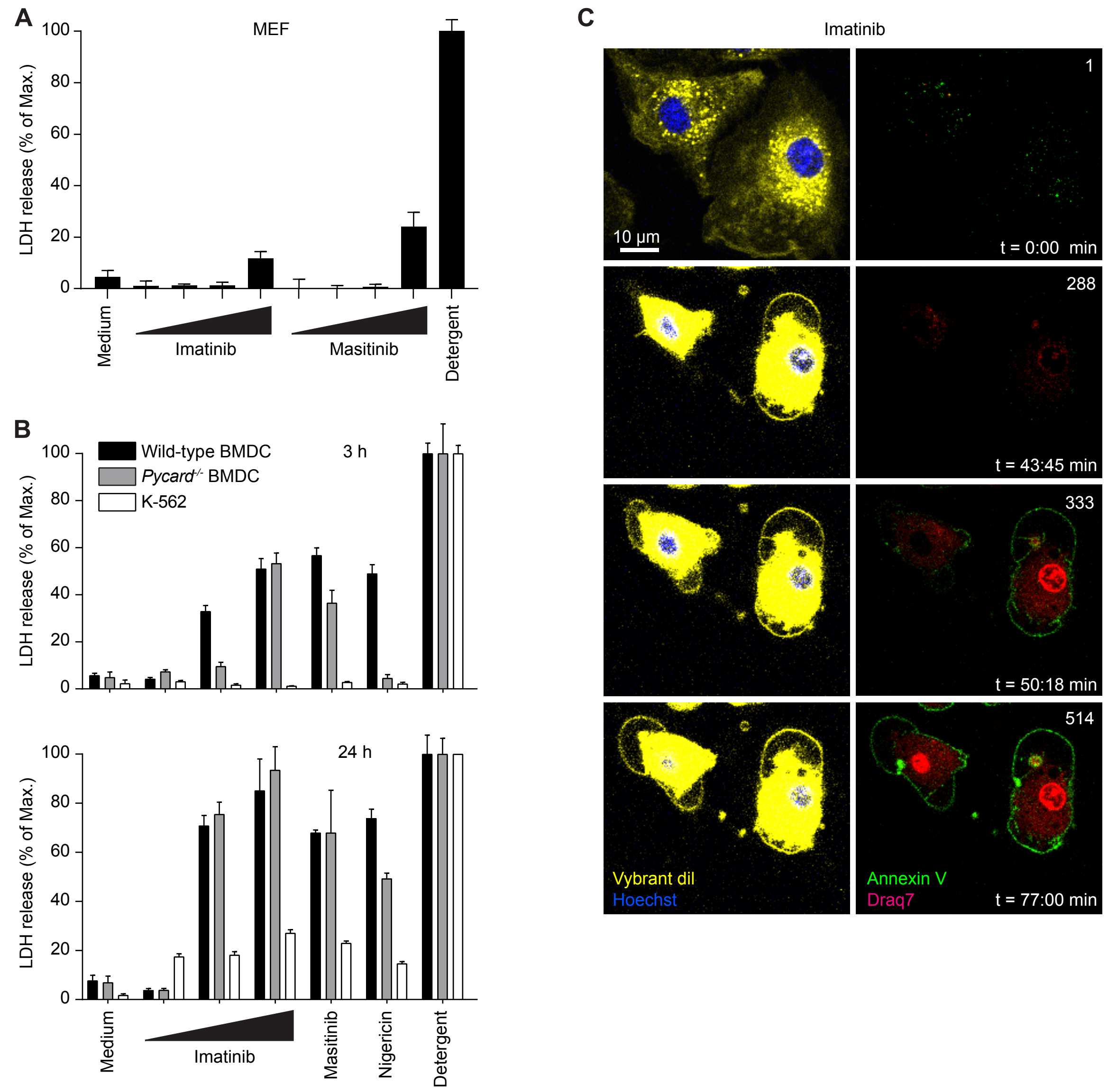


**Supplementary Fig. 5: Supplementary data corresponding to main Fig. 3: Imatinib and masitinib induce lytic death specifically in myeloid cells**

(**A**) MEFs were incubated with increasing concentrations of imatinib (10 µM, 20 µM, 40µM, 60 µM) or masitinib (5 µM, 10 µM, 20 µM, 40 µM) for 6 h. Cells were lysed as a positive control for maximum lysis ("Detergent"). LDH was determined from cell-free supernatants and is depicted as mean ±SD of technical triplicates.

(**B**) Wild-type, ASC-deficient BMDCs and K-562 cells were primed and then stimulated with imatinib (20 µM, 40 µM, 80 µM), 40 µM masitinib or 5 µM nigericin for 3 h and 24 h. Cells were lysed as a positive control for maximum lysis ("Detergent"). LDH was determined from cell-free supernatants and is depicted as mean ±SD of technical triplicates.

(**C**) LPS-primed ASC-deficient *Pycard*$^{-/-}$ BMDCs were stimulated with 60 µM imatinib or 40 µM masitinib and the cells were subsequently labelled with Annexin-V/ 7-AAD and analyzed by flow cytometry.

(**D**) LPS-primed, ASC-deficient *Pycard*$^{-/-}$ BMDMs were stained with FITC Annexin V (green), cell membrane was stained with Vybrant DiI Cell-Labeling and nuclei were localized with

Hoechst 33342 (blue) and DRAQ7 (red). The cells were then stimulated with 40 µM imatinib and live confocal imaging was performed.

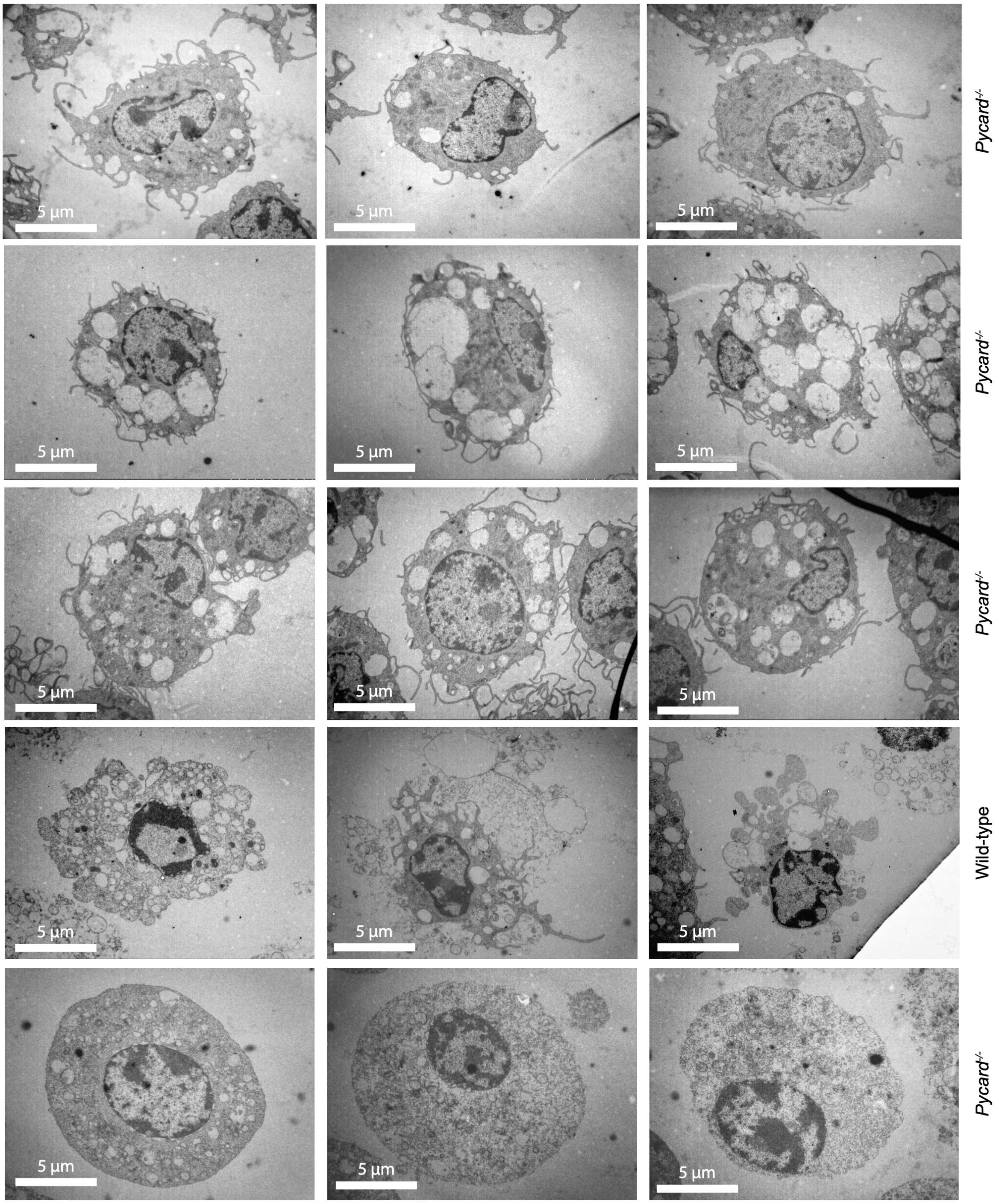


**Supplementary Fig. 6: Supplementary data corresponding to main Fig. 3: Imatinib and masitinib induced myeloid cell death is morphologically distinct from apoptosis and pyroptosis**

LPS-primed wild-type or ASC-deficient *Pycard*$^{-/-}$ BMDMs were stimulated with 40 µM imatinib, 20 µM masitinib; 5 µM nigericin, 10 µM raptinal, or DMSO for 2 h. Cells were fixed by adding 1% glutaraldehyde and analyzed by transmission electron microscopy.

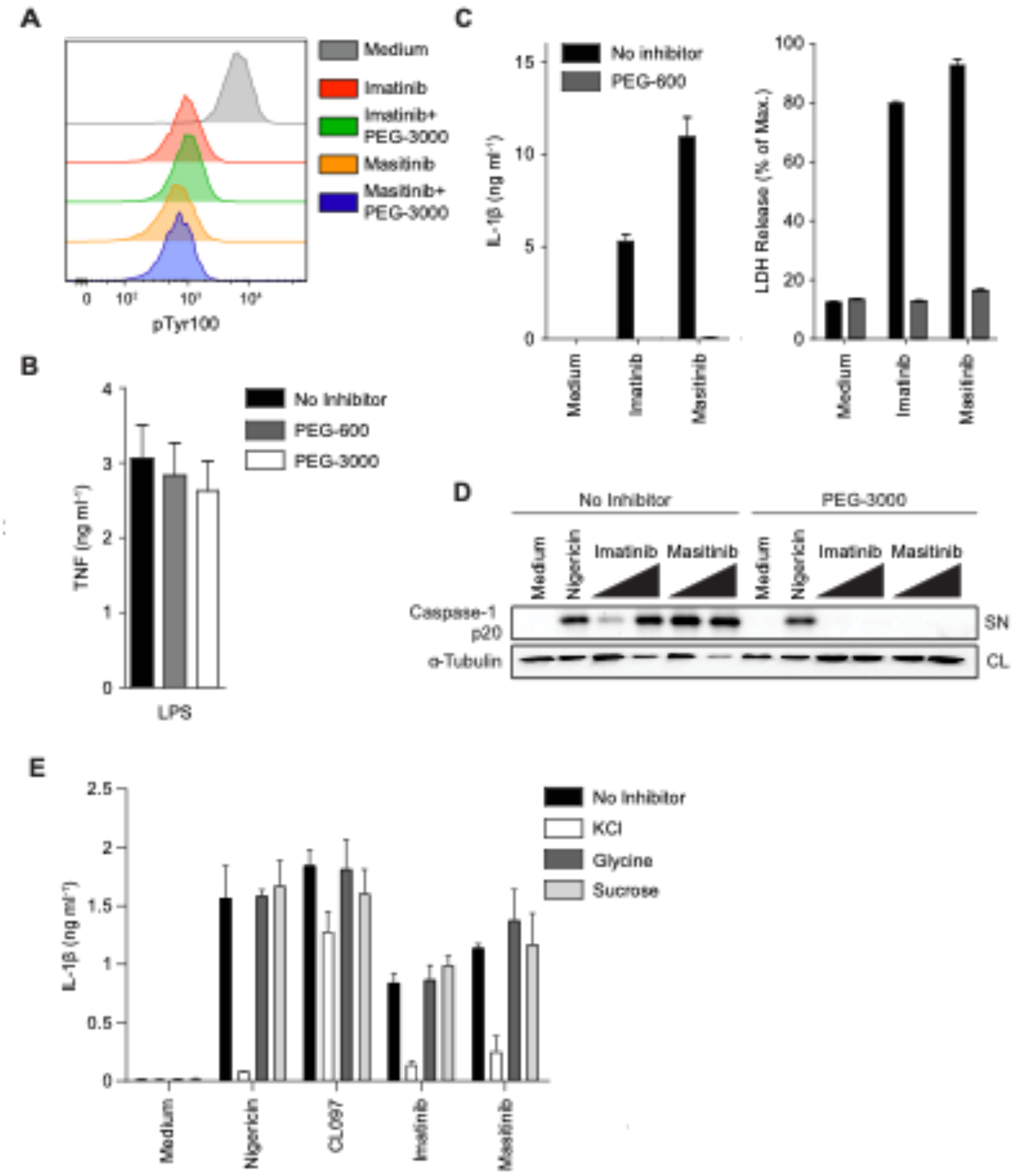


**Supplementary Fig. 7: Supplementary data corresponding to main Fig. 4: PEG rescues TKI-driven cell lysis and inflammasome activation**

(**A**) K-562 cells were incubated with 10 mM PEG-3000 or medium 30 min prior to stimulation with 40 µM imatinib or 40 µM masitinib for 3 h. Cells were stained with Zombie Aqua fixable viability dye, fixed and permeabilized, stained with phospho-tyrosine antibody and analyzed by flow cytometry.

(**B**) Corresponding to main Fig. 4D: BMDCs were left untreated or treated with 50 mM PEG-600 or 5 mM PEG-3000 and then stimulated with LPS and TNF secretion was measured.

(**C**) LPS-primed wild-type BMDCs were treated with 100 mM PEG-600 for 30 min and subsequently stimulated with 60 µM imatinib or 40 µM masitinib and IL-1β secretion and LDH release were measured.

(**D**) Immunoblot analysis of cell-free supernatants of LPS-primed BMDCs, pretreated with 5 mM PEG-3000 for 30 min and then stimulated with 5 µM nigericin or 20 µM and 40 µM of the indicated TKIs.

(**E**) LPS-primed BMDCs were incubated with 60 mM KCl, 50 mM glycine, 50 mM sucrose or left untreated for 30 min prior to stimulation with 5 µM nigericin, 100 µM CL097, 40 µM imatinib and 20 µM masitinib for 3h and IL-1β secretion was measured.

Cytokine secretion and LDH release were determined by ELISA or using a colorimetric assay, respectively, from cell-free supernatants and data are depicted as mean ±SD of technical triplicates. Results are representative of at least three independent experiments.

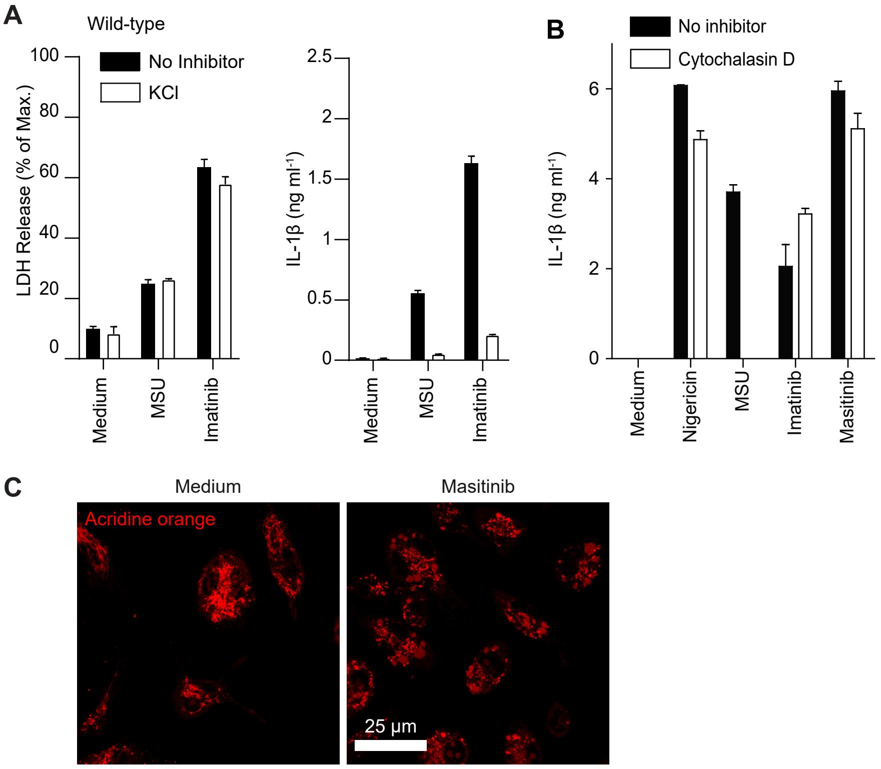


**Supplementary Fig. 8: Supplementary data corresponding to main Fig. 5: Imatinib and masitinib induce lysosomal damage**

(**A**) LPS-primed wild-type BMDCs were incubated with 50 mM KCl or left untreated for 30 min prior to stimulation with 300 µg ml$^{-1}$ MSU, 40 µM imatinib or 20 µM masitinib for 4 h and IL-1β secretion and LDH release were measured.

(**B**) LPS-primed wild-type BMDCs were treated with 3 µM cytochalasin D for 30 min or left untreated and subsequently stimulated with 5 µM nigericin, 300 µg ml$^{-1}$ MSU, 60 µM imatinib or 40 µM masitinib and IL-1β secretion was measured.

(**C**) ASC-deficient BMDMs were stained with acridine orange and incubated with 15 mM PEG-3000 30 min prior to stimulation with 20 µM masitinib. Cells were analyzed by fluorescence microscopy after 10 min.

IL-1β secretion and LDH release were determined by ELISA or using a colorimetric assay, respectively, from cell-free supernatants and data are depicted as mean ±SD of technical triplicates. Results are representative of at least three independent experiments.

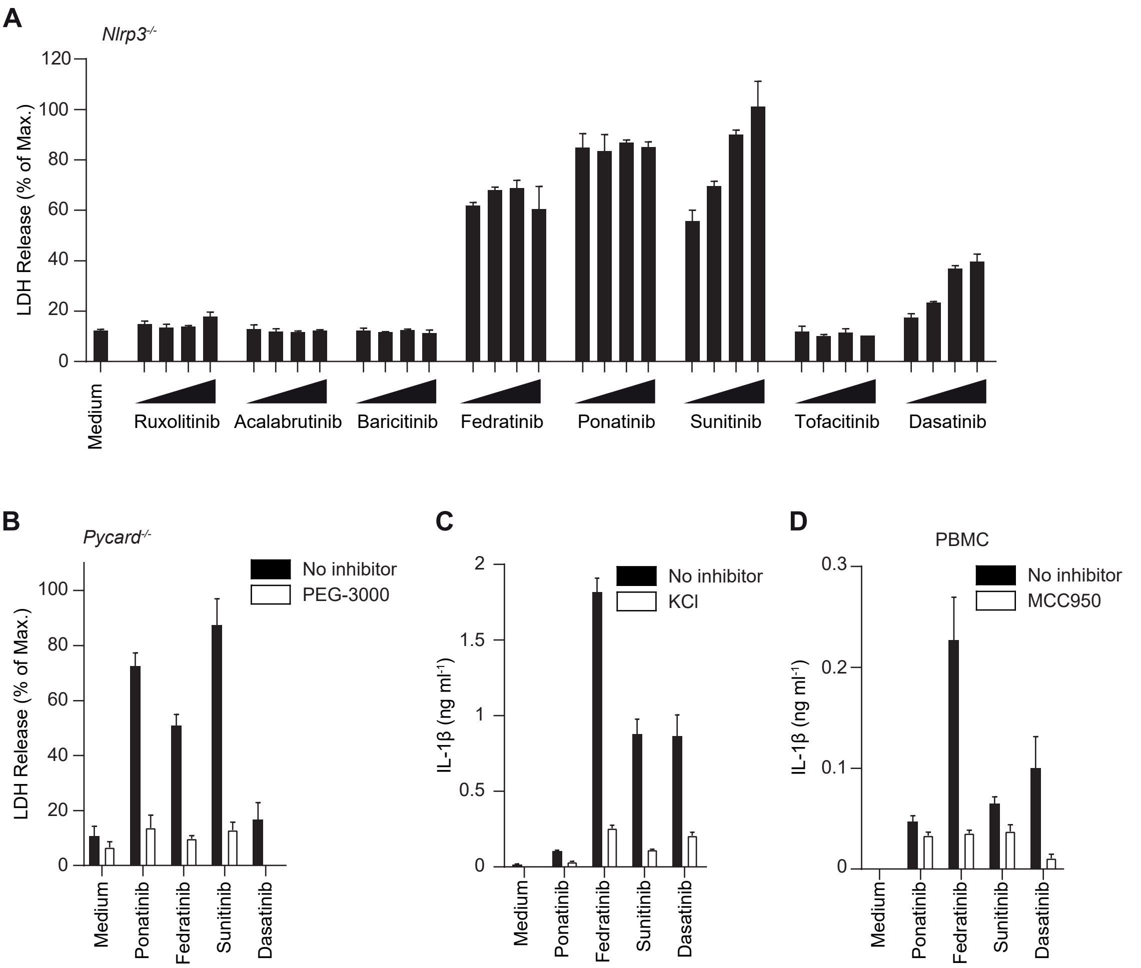


**Supplementary Fig. 9: Supplementary data corresponding to main Fig. 6: Multiple clinically relevant TKIs induce PEG-sensitive myeloid cell death and NLRP3 inflammasome activation**

(**A**) LPS-primed BMDCs from *Nlrp3*$^{-/-}$ mice were stimulated with increasing concentrations (10, 20, 40, 80 µM) of the indicated TKIs for 3 h and LDH release was measured.

(**B**) LPS-primed ASC-deficient *Pycard*$^{-/-}$ BMDCs were incubated with 15 mM PEG-3000 or left untreated for 30 min prior to stimulation with 20 µM ponatinib, 40 µM fedratinib, 60 µM sunitinib or 80 µM dasatinib for 3 h and LDH release was measured.

(**C**) LPS-primed wild-type BMDCs were treated with 60 mM KCl and stimulated with 20 µM ponatinib, 40 µM fedratinib, 40 µM sunitinib or 80 µM dasatinib and IL-1β secretion was measured.

(**D**) LPS-primed human PBMCs were treated with 3 µM MCC950 for 30 min or left untreated and subsequently stimulated with 60 µM ponatinib, 60 µM fedratinib, 40 µM sunitinib or 80 µM dasatinib for 3 h and IL-1β secretion was measured.

IL-1β secretion and LDH release were determined by ELISA or using a colorimetric assay, respectively, from cell-free supernatants and data are depicted as mean ±SD of technical triplicates. Results are representative of at least three independent experiments.

**Supplementary Table 1: Structures and trade names of the drugs analyzed throughout this study. For drugs not licensed for human use but within clinical trials, the number of trials is indicated.**

| Inhibitor | Structure |
|---|---|
| **Acalabrutinib**<br><br>Tradename:<br>Calquence™ | 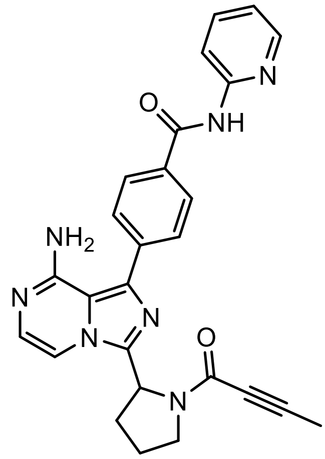 |
| **Alogliptin**<br><br>Tradename:<br>Vipidia™ | 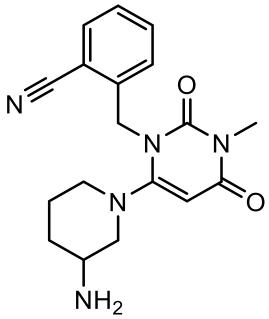 |
| **Baricitinib**<br><br>Tradename:<br>Olumiant™ | 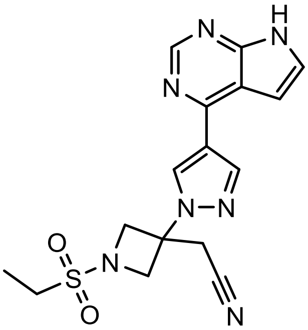 |
| **Bosutinib**<br><br>Tradename:<br>Bosulif™ | 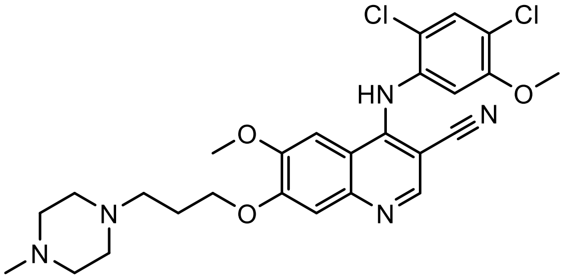 |
| **Crizotinib**<br><br>Tradename:<br>Xalkori™ | 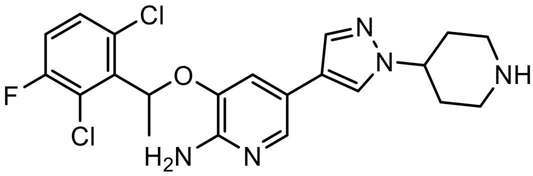 |

| | |
|---|---|
| **Dasatinib**<br><br>Tradename:<br>Sprycel™ | 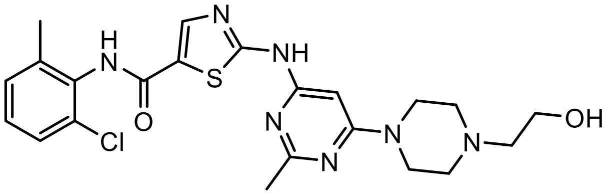 |
| **Fedratinib**<br><br>Tradename:<br>Inrebic™ | 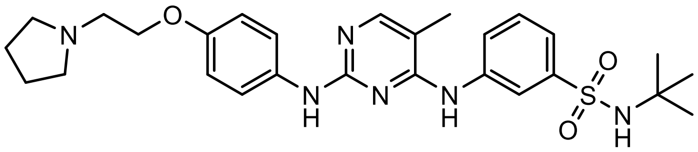 |
| **Imatinib**<br><br>Tradename:<br>Glivec™, Gleevec™ | 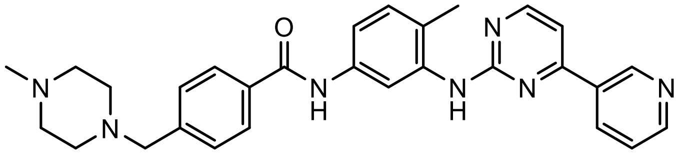 |
| **Masitinib**<br><br>Number of clinical trials: 32 | 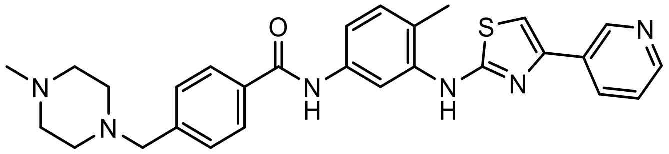 |
| **Nilotinib**<br><br>Tradename:<br>Tasigna™ | 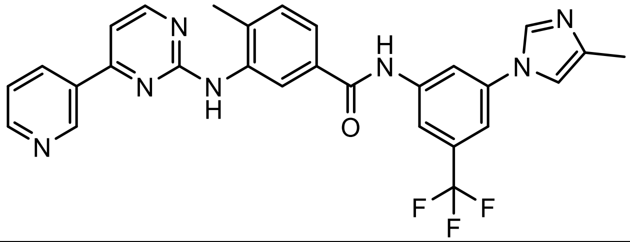 |
| **Ponatinib**<br><br>Tradename:<br>Iclusig™ | 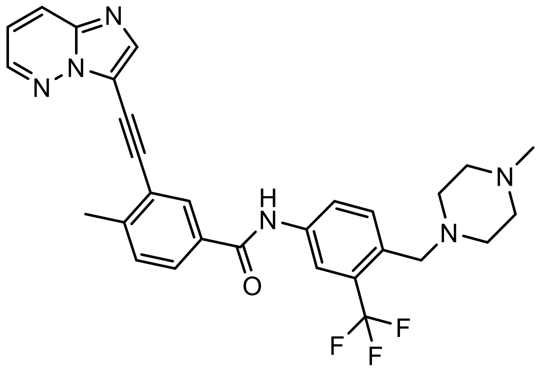 |
| **Ruxolitinib**<br><br>Tradename:<br>Jakavi™ | 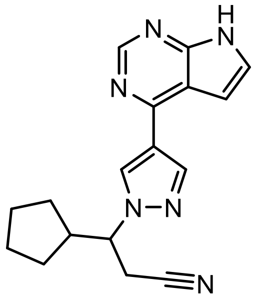 |
| **Sunitinib**<br><br>Tradename:<br>Sutent™ | 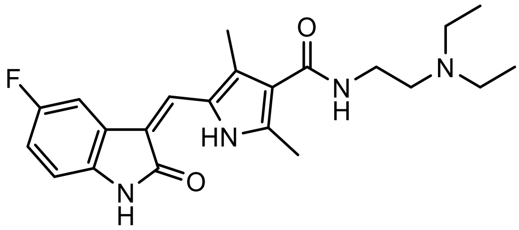 |

| | |
|---|---|
| **Tofacitinib**<br><br>Tradename:<br>Xeljanz™ | 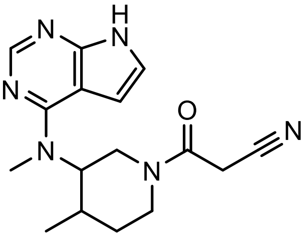 |
| **Val-boro-Pro** (VbP, talabostat, PT-1000)<br><br>Number of clinical trials: 12 | 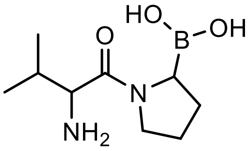 |
| **1G244** | 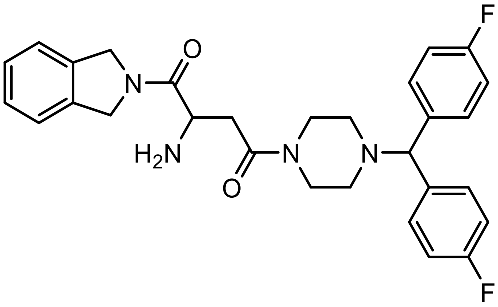 |
